# Haldane’s rule in the placenta: sex-biased misregulation of the *Kcnq1* imprinting cluster in hybrid mice

**DOI:** 10.1101/2020.05.07.082248

**Authors:** Lena Arévalo, Sarah Gardner, Polly Campbell

**Affiliations:** Department of Integrative Biology, Oklahoma State University, Stillwater, OK, USA; Department of Developmental Pathology, University Hospital, Bonn, Germany; Department of Evolution, Ecology, and Organismal Biology, University of California Riverside, Riverside, CA, USA

**Keywords:** *Ascl2*, Dobzhansky-Muller incompatibility, imprinted genes, *Mus musculus domesticus*, *Mus spretus*, parent-of-origin, *Phlda2*, postzygotic reproductive isolation, speciation

## Abstract

Mammalian hybrids often show striking asymmetries in their phenotypes both between reciprocal crosses, and between sexes in accordance with Haldane’s rule. Hybrid inviability is associated with parent-of-origin placental growth abnormalities for which misregulation of imprinted genes is a strong candidate mechanism. However, direct evidence for the involvement of abnormal imprinting and the mechanisms behind this proposed misregulation is limited. We used transcriptome and reduced representation bisulfite sequencing to evaluate the contribution of imprinted genes to a long-standing example of parent-of-origin placental growth dysplasia in the cross between the house mouse (*Mus musculus domesticus*) and the Algerian mouse (*Mus spretus*). We found little evidence for loss of imprinting and imprinted genes with biallelic expression were not misexpressed. Instead, imprinted genes with transgressive expression and methylation were concentrated in the *Kcnq1* cluster, which contains causal genes for prenatal growth abnormalities in both mice and humans. Hypermethylation of the cluster’s imprinting control region, and consequent misexpression of the genes *Phlda2* and *Ascl2*, is a strong candidate mechanism for hybrid placental undergrowth. Transgressive placental and gene regulatory phenotypes, including expression and methylation in the Kcnq1 cluster, were more extreme in hybrid males. While consistent with Haldane’s rule, male-biased defects are not expected in rodent placenta because the maternal X chromosome is effectively hemizygous in both sexes. In search of an explanation we found evidence of leaky imprinted X-chromosome inactivation in hybrid females. Supplementary expression from the paternal X-chromosome may buffer the females from the effects of X-linked incompatibilities to which males are fully exposed. Sex differences in chromatin structure on the X and sex-biased maternal effects are non-mutually exclusive alternative explanations for adherence to Haldane’s rule in hybrid placenta. The results of this study contribute to understanding of the genetic basis of hybrid inviability in mammals, and the role of imprinted genes in speciation.

## INTRODUCTION

The evolution of intrinsic postzygotic reproductive isolation in taxa with genetic sex determination is characterized by asymmetries whose direction is remarkably consistent. When one sex of hybrids is sterile or inviable it is the heterogametic sex (Haldane 1922) and the contribution of the X (or Z) chromosome to these hybrid defects is often large relative to that of the autosomes (Coyne and Orr 1989; Tao et al. 2003; Masly and Presgraves 2007; Presgraves 2018). The study of these asymmetries has revealed others: hybrid sterility typically evolves before inviability and depends on cross direction during the early stages of speciation (Coyne and Orr 2004; Turelli and Moyle 2007; Turissini et al. 2018). Across taxa, the genetic architecture of hybrid defects is largely consistent with the Dobzhansky-Muller model for the evolution of intrinsic incompatibilities, in which independently evolving loci interact negatively when combined in hybrids (Bateson 1909; Dobzhansky 1937; Muller 1942; Coyne and Orr 2004). Strikingly, loci that cause or contribute to Dobzhansky-Muller incompatibilities (DMIs) are often implicated in antagonistic coevolution within species (Presgraves 2010; McDermott and Noor 2010; Crespi and Nosil 2013; Patten 2018; Zanders and Unckless 2019), itself a consequence of asymmetries in the fitness optima of interacting partners (e.g. selfish genetic elements and host genomes, males and females, parasites and hosts).

In mammals, hybrid inviability is associated with another pronounced asymmetry: growth abnormalities that depend on cross direction are documented in four mammalian orders with the most detailed genetic studies conducted in rodents (Gray 1972; Zechner et al. 1996; Vrana et al. 1998; Brekke and Good 2014). For example, the cross between a female oldfield mouse (*Peromyscus polionotus*) and a male deer mouse (*P. maniculatus*) produces hybrid conceptuses (embryo+placenta) that are grossly over-sized and rarely survive to term. Offspring from the reciprocal cross are viable but significantly smaller than both parental species (Rogers and Dawson 1970; Vrana et al. 1998). Likewise, in the cross between the house mouse, *Mus musculus domesticus*, and the Algerian mouse, *M. spretus*, hybrid placentas are oversized (hyperplastic) when the paternal species is *M. m. domesticus* and undersized (hypoplastic) when *M. spretus* is the father (Zechner et al. 1996). Similar patterns of hybrid over- and under-growth result from reciprocal crosses in dwarf hamsters (*Phodopus*) (Brekke and Good 2014).

Imprinted genes (IGs), a small but developmentally critical group of autosomal genes, are *a priori* candidates for these hybrid growth asymmetries. IGs are highly expressed in placenta and brain, and a subset are essential regulators of placental and embryonic growth (Ferguson-Smith 2011; Barlow and Bartolomei 2014). IG inheritance is Mendelian but expression is not. Paternally expressed IGs (Pegs) are expressed predominantly or exclusively from the paternally inherited allele. Maternally expressed IGs (Megs) have the opposite expression pattern. Thus, like the X chromosome in heterogametic males, IGs are effectively hemizygous, making them strong candidate contributors to DMIs in hybrids of both sexes (Vrana et al. 2000; Vrana 2007).

The dominant hypothesis for the evolution of imprinted expression is based on parental conflict over maternal investment due to relatedness asymmetries between mothers and offspring (always 0.5) and fathers and offspring (0.5 or 0) in polyandrous mating systems (Moore and Haig 1991; Haig 2000). This adds to the appeal of IGs as speciation genes; hybrid inviability as a byproduct of antagonistic coevolution within species would be consistent with the proposition that conflict is a common driver of reproductive isolation (Presgraves 2010; Crespi and Nosil 2013; Patten 2018).

The strongest empirical support for the role of IGs in speciation comes from the *Peromyscus* cross, in which severe hybrid conceptus overgrowth is accompanied by widespread loss of imprinting (biallelic expression) (Vrana et al. 2000). The imprinted gene, *Peg3*, is a strong candidate for the autosomal locus in an X-autosome DMI for hybrid growth hyperplasia (Loschiavo et al. 2007). In *Mus* hybrids, several imprinted genes have biallelic expression and altered methylation in some tissues (Shi et al. 2005), but candidate gene studies did not establish a strong connection between IGs and placental growth phenotypes (Zechner et al. 2002; 2004). Indeed, the genetic architecture of hybrid growth dysplasia in *Mus* exhibits the same fundamental asymmetries that characterize intrinsic postzygotic reproductive isolation in general: hybrid male placental phenotypes are more extreme than those of hybrid females and the effect of X chromosome genotype is larger in males (Zechner et al. 1996; Hemberger et al. 1999; 2001). Because the paternally inherited X chromosome is normally imprinted (silenced) in female mouse placenta (Tagaki and Sasaki 1975), both sexes are exposed to recessive-acting DMIs on the X. Adherence to Haldane’s rule in the placenta is therefore surprising.

Here, we present the first genome-scale analysis of this long-standing example of parent-of-origin-dependent hybrid growth asymmetries. We use placental morphology (weight and histology), placental transcriptome and reduced representation bisulfite sequencing to characterize the relationships between placental hypoplasia, gene expression, DNA methylation, and sex-specific effects in the cross between female *M. m. domesticus* and male *M. spretus*. Placental transcriptome data were included in Arévalo and Campbell (2020) but sex differences in expression and analyses of allele specific expression are reported here for the first time. All other data and analyses are new to this study. We address three main questions. 1) Are imprinted genes strong candidates for hybrid placental abnormalities? We use two types of evidence to evaluate this question: first, transgressive expression of imprinted genes themselves and second, functional associations between transgressively expressed genes (imprinted or not) and imprinted genes that retain allele specific (effectively hemizygous) expression in hybrid placenta. 2) Does sex-specific expression and methylation recapitulate placental phenotypes? Specifically, are there more genes with transgressive expression and/or methylation in hybrid males vs. females? 3) Are there sex differences in X-linked expression that could underlie the more extreme placental phenotype in hybrid males? To address this question we tested for sex-specific overrepresentation of transgressive expression on the X chromosome and evaluated the possibility that leaky paternal X chromosome inactivation buffers female hybrids from potential X-linked incompatibilities that are exposed in males.

## METHODS

### Animals, tissue collection, and phenotypic analyses

Mice used in this study were maintained on a 12:12 light:dark cycle with lights on at 9:00 AM and were provided with 5001 Rodent Diet (LabDiet, Brentwood, MO, U.S.A.) and water ad lib. All animal procedures were approved by the Oklahoma State University IACUC under protocol #141-AS. *M. m. domesticus* (hereafter, *Dom*) was represented by the wild-derived inbred strain WSB/EiJ (Jackson Laboratory) and *M. spretus* (hereafter, *Spret*) was represented by the wild-derived inbred strain SFM/Pas (Montpellier Wild Mice Genetic Repository). We conducted three crosses (female listed first): *Dom* X *Dom*, *Dom* X *Spret*, *Spret* X *Spret*. Prior to pairing, females were placed in a cage with soiled conspecific male bedding for ~48 hrs. to induce receptivity to mating (Whitten 1956). Mice were paired between 5:00 and 6:00 PM, left undisturbed for two nights, and split on the morning of the second day. The second night was counted as embryonic day 0 (e0). Pregnant females (n=5/cross) were euthanized by cervical dislocation between 10:00 and 11:00 AM on embryonic day 17-18 (e17.5). Placentas were separated from embryos, and the maternally-derived decidual layer was removed as described (Qu et al. 2014). Embryos and placentas were weighed and transferred immediately to RNAlater (Thermo Fisher, USA), kept at 4°C overnight to allow RNAlater perfusion, and stored at −20°C until RNA extraction. Since litter size can affect placenta and embryo size (Ishikawa et al. 2006) we corrected weights for litter size (individual placenta or embryo weight * litter size) prior to analysis.

Placentas used for histology were fresh frozen and stored at −80°C until cryosectioning. Frozen placenta was cryosectioned at 20μm on a Leica CM 1950 cryostat and mounted on Fisher Superfrost Plus slides. Mounted sections were immediately stained with hematoxylin and eosin using the Thermo Scientific Shandon Rapid-Chrome H & E Frozen Section Staining Kit (Thermo Fisher). Stained sections were imaged at 10x with a Leica DM IL LED microscope, equipped with a QICAM Fast 1394 camera with QCapture software.

### Sample preparation and sequencing

Embryo sex was determined by PCR for the Y-linked gene, *Zfy1*. Total RNA was extracted from one male and one female placenta per litter (n=5 males/cross; n=4 females [hybrid cross] and n=5 females [conspecific cross]) using the AllPrep RNA/DNA Mini Kit (Qiagen), and was stored at −80°C until sequencing. Illumina library preparation was done at the sequencing facility (Novogene, Sacramento, CA) using the NEB Next Ultra RNA library prep kit for Illumina. Individual libraries were sequenced on the Illumina HiSeq 4000 platform, producing >30 million, 150bp paired-end reads per sample.

### Transcriptome data processing and analysis

QC, trimming of raw sequencing reads, filtering and mapping were performed as described (Arévalo and Campbell 2020). To improve comparability, all samples (*Dom, Spret* and hybrid) were mapped to a pseudo-hybrid genome, generated using the genome preparation tool of the program SNPsplit (Krueger and Andrews 2016). Briefly, SNPs from both *Dom* (WSB/EiJ) and *Spret* (SPRET/EiJ) relative to the mouse genome (GRCm38.89) available from the Ensembl FTP server (ftp://ftp.ensembl.org) were introduced into the mouse genome. SNPs between *Dom* and *Spret* were then N-masked to allow mapping of both *Dom*- and *Spret*-derived reads. To account for possible mapping bias against *Spret*, we randomly downsampled all alignment files to ~40 million reads using SAMtools 0.1.19 (Li et al. 2009). For analysis of allele specific expression (ASE) in hybrids we used SNPsplit to split the hybrid alignment files, separating reads originating from the *Spret* and *Dom* alleles. Mapped reads were assigned to the *Dom* or *Spret* allele based on a list of SNPs between the WSB/EiJ and SPRET/EiJ, and the overlap with N-masked positions in the mouse genome.

Transcript quantification and annotation was done using StringTie 1.3.3 (Pertea et al. 2015). Mouse genome annotation information (GRCm38.89) was retrieved from the Ensembl FTP server (ftp://ftp.ensembl.org). We used the python script (preDE.py) included in the StringTie package to prepare gene-level count matrices for analysis of differential expression (DE) and ASE in DESeq2 1.16.1 (Love et al. 2014). Pseudogenes were removed from the count matrices based on “biotype” annotation information extracted from Biomart (R-package biomaRt, Durinck et al. 2005). Low counts were removed by the independent filtering process implemented in DESeq2 (Bourgon et al. 2010). The adjusted p-value (Benjamini-Hochberg method) cutoff for DE was set at 0.05. The log2 fold change cutoff was set to > 0.5 for standard DE analysis and to >1 for ASE analysis. Normalized DESeq2 read counts for ASE were used to calculate the proportion of reads expressed from the *Dom* allele.

### Reduced representation bisulfite sequencing data processing and analysis

We used genome-wide reduced representation bisulfite sequencing (RRBS) to identify differentially methylated regions in CpG islands. CpG enriching restriction digest, bisulfite treatment, size selection, library preparation and sequencing were performed by the sequencing facility (NXT-Dx, Ghent, Belgium). Sequencing was performed on the Illumina HiSeq 4000 platform, producing >13 million, 50bp paired-end reads per sample.

QC of raw sequencing reads and trimming were performed in Trim Galore! 0.4.5 (Babraham Bioinformatics, http://www.bioinformatics.babraham.ac.uk/projects/trim_galore), using a phred score cutoff of 20 and minimum sequence length of 20 after trimming. The pseudo-hybrid genome was bisulfite converted and indexed for mapping using the Bismark-genome-preparation tool and the RRBS reads were mapped to the converted pseudo-hybrid genome using the Bismark Bisulfite Aligner (Babraham Bioinformatics, http://www.bioinformatics.babraham.ac.uk/projects/bismark/). We filtered the resulting alignment files using SAMtools 0.1.19 (Li et al. 2009), retaining only high quality (MAPQ score 40), uniquely mapped, paired reads for analysis. To account for possible mapping bias against *Spret*, we randomly downsampled all alignment files to ~10 million reads using SAMtools 0.1.19 (Li et al. 2009). To analyze allele specific methylation (ASM) in hybrids we split the hybrid alignment files as described above for RNAseq alignments.

CpG methylation information was extracted from the alignment files using the Bismark-methylation-extractor tool (Babraham Bioinformatics, http://www.bioinformatics.babraham.ac.uk/projects/bismark/). Differential methylation (DM) and ASM were analyzed in SeqMonk (Babraham Bioinformatics, http://www.bioinformatics.babraham.ac.uk/projects/seqmonk/). CpG methylation percentage of CpG islands was determined using the bisulfite feature methylation pipeline in SeqMonk. DM and ASM were determined for CpG islands located in gene promoter regions using logistic regression with Benjamini-Hochberg correction. Log2 fold change between compared samples was calculated and cutoff was set to > 0.5 for standard DM analysis and to >1 for ASM analysis. CpG islands in and around imprinted genes, including imprinting control regions (ICRs), were tested for DM and ASM in the same manner.

### Gene ontology (GO) term and pathway over-representation analysis

We performed GO term and pathway over-representation analyses on relevant lists of genes from DE, ASE, and DM analyses using the PANTHER gene list analysis tool with Fisher’s exact test and FDR correction (Mi et al. 2017). We tested for over-representation based on the GO annotation database (Biological Processes) (released 07-Jan-2017, Ashburner et al. 2000; The Gene Ontology Consortium 2017) and the Reactome pathway database (version 58, Fabregat et al. 2017).

### X-linked and imprinted gene network analyses

We used String (v11, Szklarczyk et al. 2019) to construct gene networks for sets of genes that are candidate effectors of hybrid placental abnormalities in one or both sexes: X-linked genes with significant expression from the paternal allele in hybrid females and placental imprinted genes with statistically significant allelic expression bias in hybrids. Whereas X-linked expression in female mouse placenta is predominantly maternal, genes with significant expression from the paternal allele might buffer hybrid females from recessive-acting incompatibilities to which hybrid males are fully exposed. Canonically expressed imprinted genes are functionally hemizygous, and are therefore *a priori* candidates for contribution to hybrid incompatibilities in both sexes. Parameters for string were set to a maximum of 20 interactors in the first shell and a maximum of 5 interactors in the second shell, with a minimum required interaction score of 0.4. Results from text-mining, databases, experiments, and gene fusions were considered. We tested for overrepresentation of transgressively-expressed genes in both X-linked and IG gene networks using Fisher’s exact test.

## RESULTS

### Reduced viability in hybrid litters

There was a significant effect of litter genotype on embryonic viability in late gestation that was due to a lower proportion of viable embryos in hybrid litters (One-way ANOVA: F(2)=5.41, p=0.012, Tukey HSD: Hybrid-*Dom*: diff=0.15, p=0.041, Hybrid-*Spret*: diff=0.18, p=0.012, *Dom-Spret*: diff=0.028, p=0.86) (Table 1). Inviable conceptuses were consistently dark-colored and highly condensed with no discernable distinction between embryonic and extraembryonic tissues, phenotypes indicative of late stage of resorbtion at e17.5 and thus early lethality (Pang et al. 2014). Litter size, counting only viable embryos, was slightly smaller for hybrid relative to *Dom* and *Spret* litters but the effect of genotype was not significant (One-way ANOVA: F(2)=0.51, p=0.61) (Table 1). There were numerically more inviable hybrid males (*n*=6) than females (*n*=2) but within-litter sex ratios for viable embryos were also male-biased in hybrids (mean proportion males, 0.57 ± SD 0.27).

**Table 1.** Litter size and viability.

|  | *n* litters | Litters with<br>inviable<br>embryos | Total<br>inviable<br>(sex) | Mean proportion<br>viable embryos<br>(SD) | Mean<br>litter size<br>(SD) |
| --- | --- | --- | --- | --- | --- |
| *Dom x Dom* | 9 | 1 | 2 (f) | 0.96 (0.13) | 5.0 (1.1) |
| *Spret x Spret* | 10 | 1 | 1 (m) | 0.98 (0.05) | 5.2 (1.6) |
| *Dom x Spret* | 7 | 5 | 2 (f), 6 (m) | 0.80 (0.16) | 4.6 (1.0) |

### Hybrid placental undergrowth is more extreme in males

Prior work on the *Dom* x *Spret* cross reported a hypoplastic (undersized) placenta in F1 hybrids relative to the maternal species (*Dom*) but did not test for hypoplasia relative to *Spret* (Zechner et al. 1996). We found that both male and female hybrid placentas weighed significantly less than placentas of both parental species with no significant differences between parental species (One-way ANOVA, females (*n*=36): F(2)=13.16, p<0.001, Tukey HSD: Hybrid-*Dom* (*n*=24): diff=0.08, p<0.001, Hybrid-*Spret* (*n*=22): diff=0.07, p<0.001, *Dom-Spret* (*n*=26): diff=0.005, p=0.93; One-way ANOVA, males (*n*=40): F(2)=54.79, p<0.001, Tukey HSD: Hybrid-*Dom* (*n*=25): diff=0.1, p<0.001, Hybrid-*Spret* (*n*=28): diff=0.1, p<0.001, *Dom-Spret* (*n*=27). diff=0.008, p=0.82) (Fig. 1). Hybrid male placentas weighed significantly less than hybrid female placentas (One-way ANOVA: F(1)=5.994, p=0.02, *n*=23). There was no significant difference between sexes for *Dom* or *Spret* (One-way ANOVA: *Dom* (*n*=26): F(1)=4.063, p=0.06, *Spret* (*n*=27): F(1)=0.537, p=0.47) (Fig. 1). Embryo weights did not differ significantly in any comparison (data not shown).

**Figure 1.**
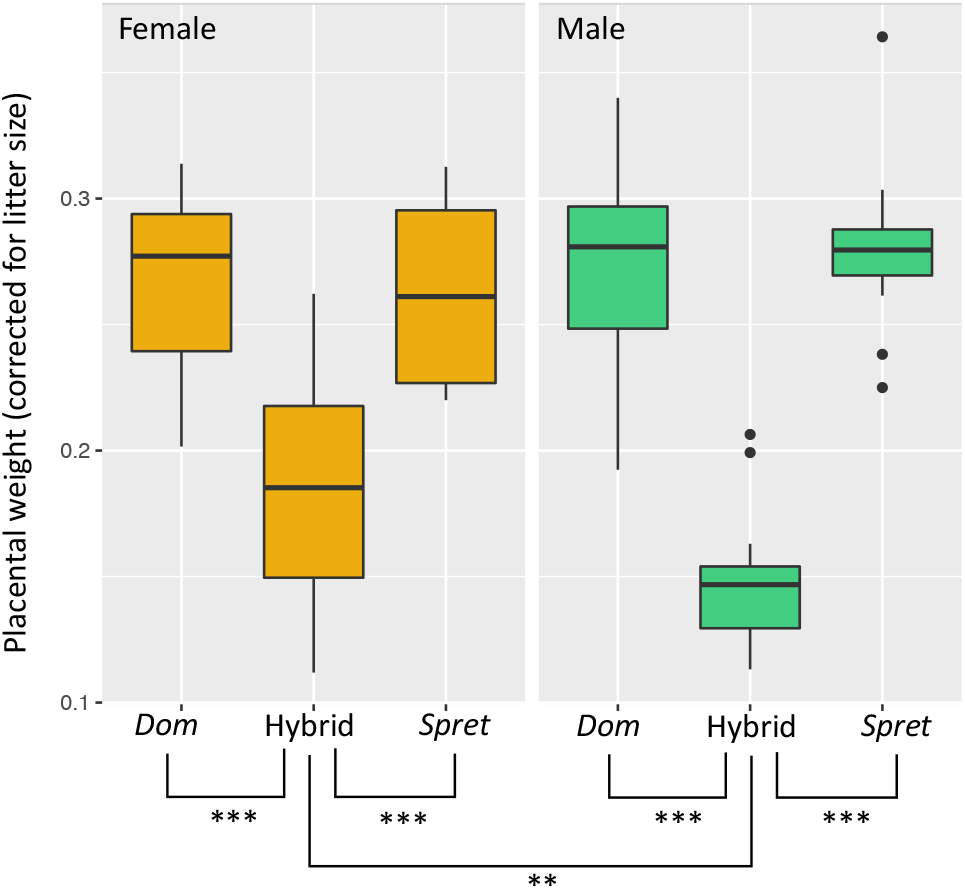
Sex-specific placental weight in hybrids and parental species, *Mus m. domesticus* (*Dom*) and *M. spretus* (*Spret*). Boxplots show litter size-corrected (placental weight [g] * litter size) placental weights for females (gold) and males (green). Asterisks indicate significance of two-way ANOVA followed by Tukey HSD post-hoc test (** *p* < 0.01, *** *p* < 0.001).

Placental hypoplasia in this cross was proposed to be due in part to a reduced spongiotrophoblast layer (Zechner et al. 1996), which lies at the maternal-fetal interface in the junctional zone (John 2013). While we did not differentiate placental cell types, placental histology confirmed that the junctional zone of the hybrid placenta was qualitatively narrower relative to that of both parental species (Fig. S1).

### Differential expression in the hybrid placenta

We tested for differential placental expression in three pairwise comparisons per sex: hybrid vs. *Dom*, hybrid vs. *Spret*, and *Dom* vs. *Spret*. We considered genes with log2 fold change (LFC) of expression ≥ 0.5 (1.5 times higher or lower expression) and Benjamini-Hochberg-corrected p≤0.05 as significantly differentially expressed (DE). Hybrid gene expression that is significantly higher or lower compared to both parental species is defined as transgressive.

In hybrid male placentas, 9.21% of all tested genes were expressed higher and 7.21% lower compared to *Dom* male placentas (up: 1,625/17,639, including 6 IGs; down: 1,271/17,639, including 7 IGs), and 14% of genes were expressed higher and 8.86% lower compared to *Spret* male placentas (up: 2,507/17,902, including 13 IGs; down: 1,587/17,902, including 9 IGs) (Fig. 2A and Dataset S1). In hybrid female placentas, 6.38% of all tested genes were expressed higher and 6.10% lower compared to *Dom* female placentas (up: 1,113/17,433, including 9 IGs; down: 1,064/17,433, including 2 IGs), and 12.08% of genes were expressed higher and 7.78% lower compared to *Spret* female placentas (up: 2,137/17,691, including 16 IGs; down: 1,377/17,691, including 7 IGs) (Fig. 2A and Dataset S1).

**Figure 2.**
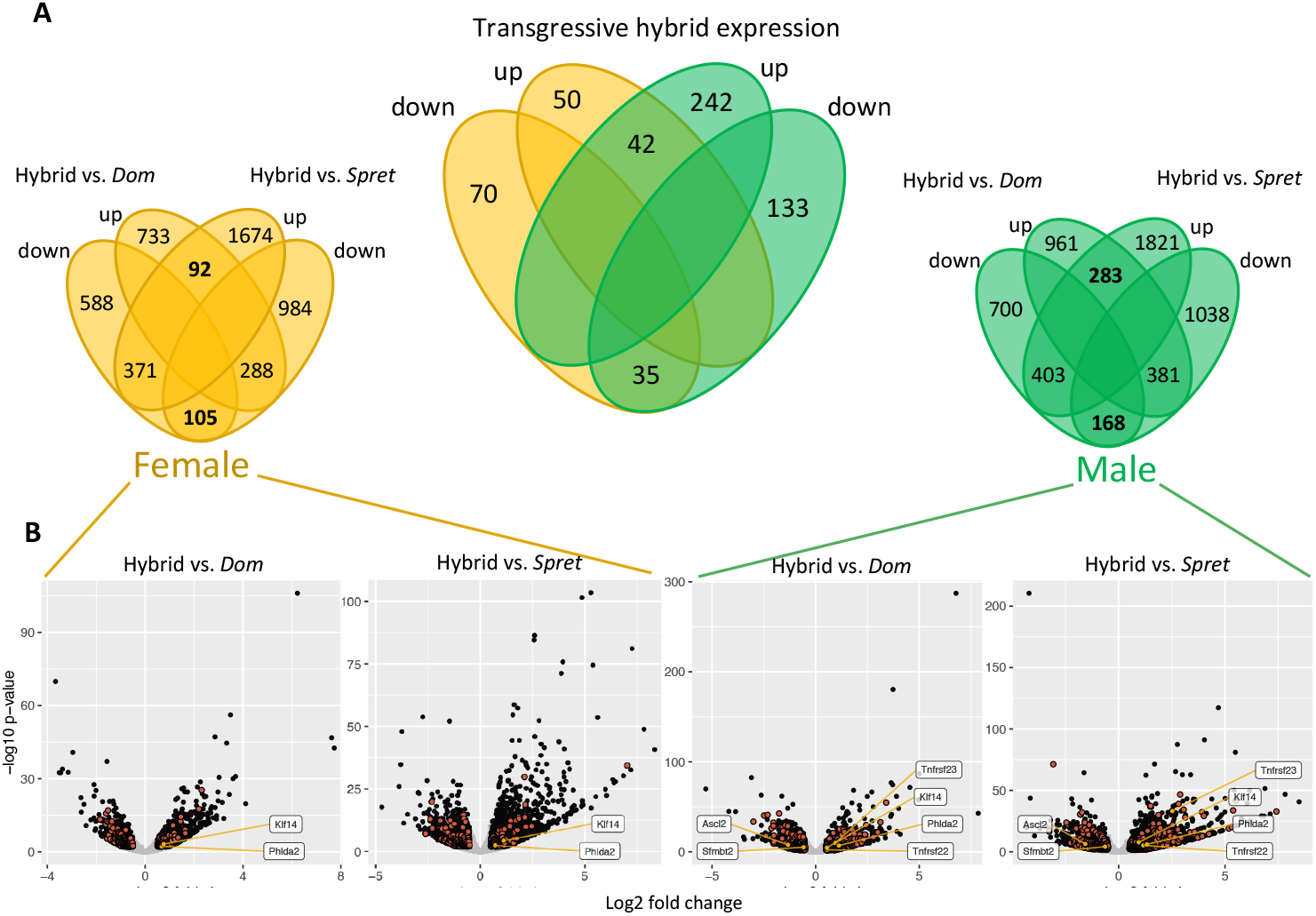
Differential expression (DE) in female (gold) and male (green) hybrid placenta relative to *Mus m. domesticus* (*Dom*), and *M. spretus* (*Spret*). A) Small Venn diagrams show the overlap of DE genes between Hybrid vs. *Dom* and Hybrid vs. *Spret*; up = genes expressed higher and down = genes expressed lower in hybrids compared to parental species; transgressive expression is indicated in bold. The large Venn diagram shows the overlap of transgressively expressed genes between female and male hybrids. B) Volcano plots of DE in Hybrid vs. *Dom* and Hybrid vs. *Spret* placentas for females and males. Significantly DE genes (FDR ≤ 0.05 and log2 fold change ≥ 0.5) are shown in black, transgressively expressed genes in red; transgressively expressed IGs (orange) are labeled.

In hybrid male placentas 283 genes were expressed transgressively higher and 168 transgressively lower. In hybrid females 92 genes were expressed transgressively higher and 105 transgressively lower (Fig. 2A). Of these genes, 42 with transgressively higher and 35 with transgressively lower expression were common to both sexes. Transgressively down-regulated genes were enriched for prolactin receptor signaling in both sexes with additional enrichment in females (Dataset S1). Transgressively expressed genes were enriched on the X chromosome in both sexes (Fisher’s exact test: Males: p=0.005, odds ratio = 1.79, Females: p=0.001, odds ratio = 2.40; Fig. S2 and Dataset S1). While there was more transgressive X-linked expression in males (29 genes) vs. females (17 genes), enrichment relative to the autosomes was higher in females (Fig. S2 and Dataset S1).

### Local, not global, misregulation of imprinted genes in the hybrid placenta

In hybrid male placentas 4 IGs (*Tnfrsf22, Tnfrsf23, Phlda2* and *Klf14*) were transgressively upregulated and two, (*Ascl2* and *Sfmbt2*) were transgressively down-regulated. *Tspan32* was significantly DE compared to both parental species but intermediate between the two. Five of these misexpressed IGs belong to the same imprinting cluster (IC2) on the distal part of mouse chromosome 7 (dist7), and are normally maternally expressed (Table 2 and Dataset S1). In hybrid female placentas 2 IGs (*Phlda2* and *Klf14*) were transgressively upregulated and *Th* was significantly DE and intermediate between the two parental species (Table 2 and Dataset S1).

**Table 2.**
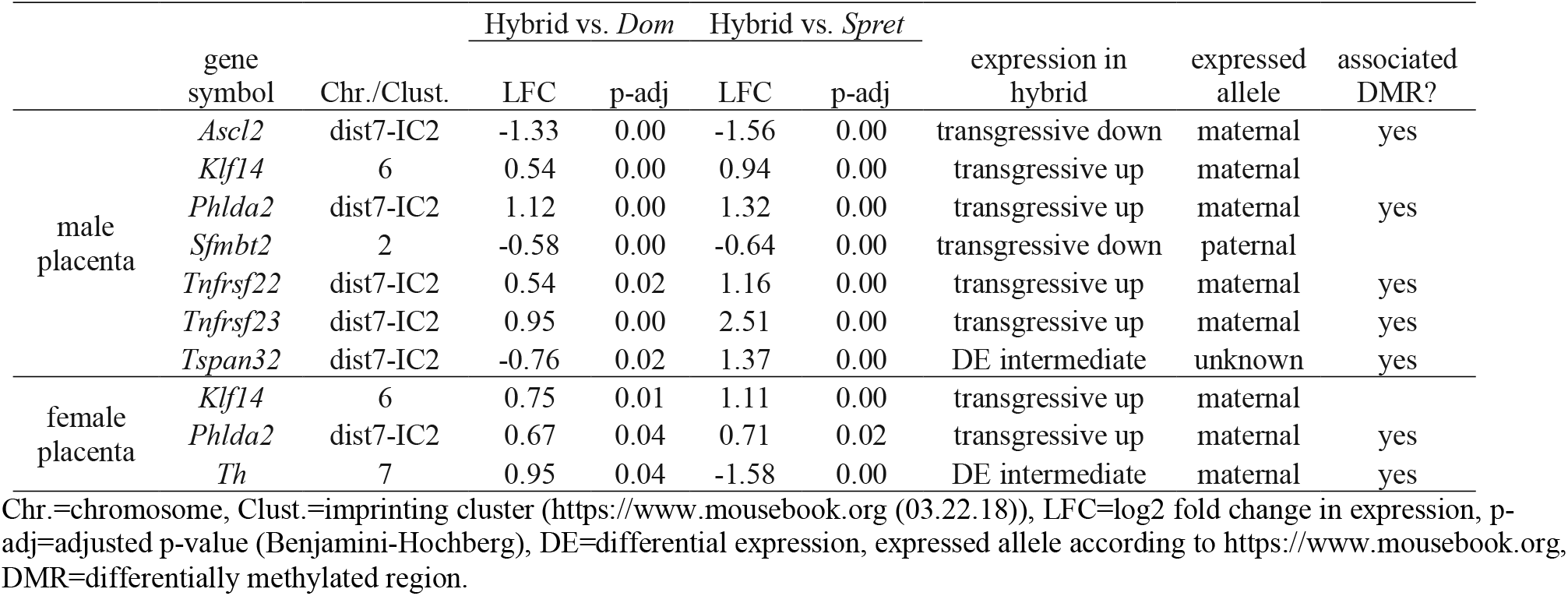
Differential expression of imprinted genes in male and female hybrids compared to both parental species.

### Global allelic expression patterns in the placenta

In both male and female hybrid placentas, autosomal allelic expression patterns were assessed by calculating the proportion of reads expressed from the *Dom* allele. The mean was very close to 0.5 for both sexes, as expected (male placenta: 0.529, female placenta: 0.527). However, the distribution was slightly skewed to the right, towards a higher proportion of *Spret* reads for both sexes (Skewness: male placenta: 0.441, female placenta: 0.474) (Fig. 3, Dataset S2).

**Figure 3.**
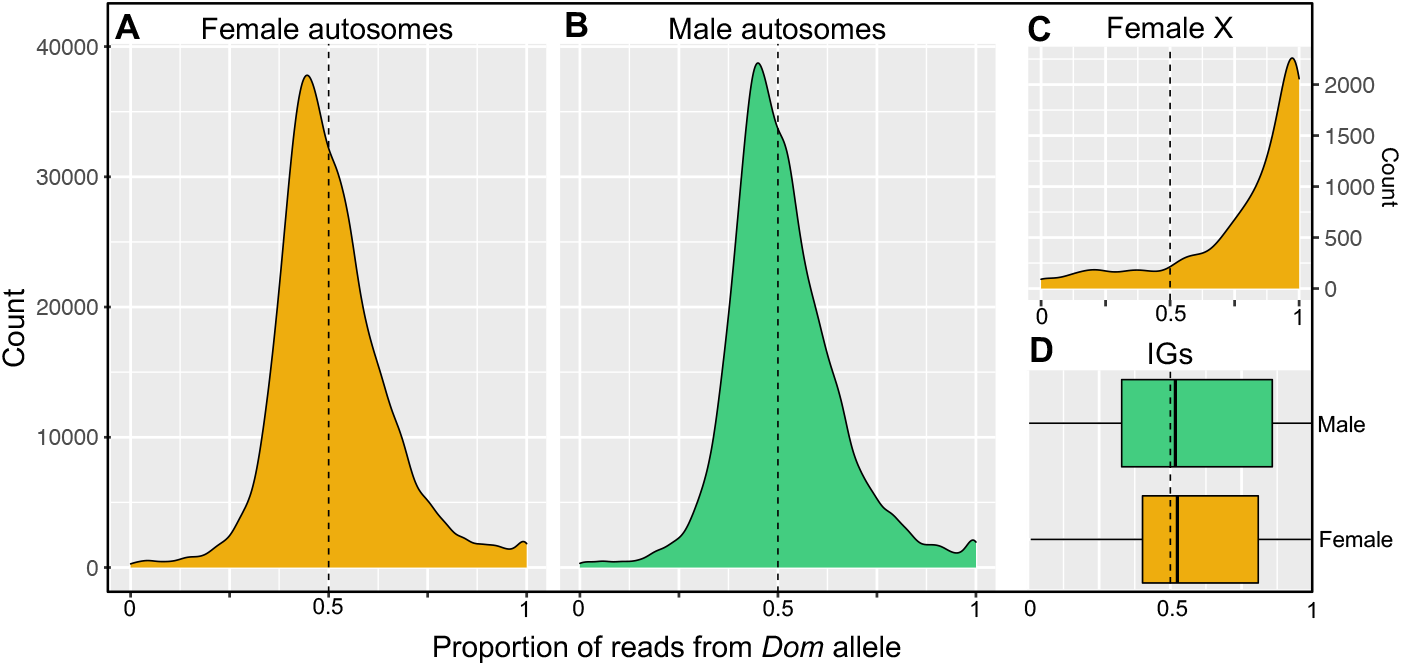
Patterns of allele specific expression in female (gold) and male (green) hybrid placenta. Large density plots show the distribution of the proportion of reads expressed from the *Dom* allele for autosomal genes in (A) females and (B) males. (C) Density plot showing the proportion of reads expressed from the *Dom* allele for the X-chromosome in females. (D) Box plot showing the proportion of reads expressed from the *Dom* allele for imprinted genes in both sexes.

### Allele specific expression in the placenta

We evaluated allele specific expression (ASE) in male and female hybrid placentas by testing for DE between alleles. We considered genes with log2 fold change (LFC) of expression ≥ 1 (2 times higher or lower expression from one allele compared to the other), and Benjamini-Hochberg-corrected p ≤ 0.05, as showing significant allele specific expression (DE). To test for possible loss of imprinting, imprinted genes with LFC ≤ 0.3 and Benjamini-Hochberg-corrected p ≥ 0.1 were considered biallelically expressed. Only the subset of genes with robust evidence for imprinting in e17.5 mouse placenta (n=36; Babak et al. 2015) were considered.

In hybrid male placentas, 12.67% and 5.54% of all tested genes had significantly higher expression from the *Dom* or *Spret* allele, respectively (*Dom* ASE: 2,019/16,127, including 25 IGs; *Spret* ASE: 894/17,639, including 17 IGs). Two of the 36 IGs evaluated met our criteria for biallelic expression: *Copg2*, and *Gnas* (Dataset S2, Fig. S3). In hybrid female placentas, 12.52% and 4.52% of all tested genes had significantly higher expression from the *Dom* or *Spret* allele, respectively (*Dom* ASE: 2,010/15,853, including 21 IGs; *Spret* ASE: 717/15,853, including 11 IGs). Three IGs were biallelically expressed: *Zdbf2, Copg2*, and *Gnas* (Dataset S2, Fig. S3).

As expected, X chromosome-wide allelic expression in hybrid female placenta was highly skewed towards the *Dom* allele (Mean: 0.783, Skewness: −1.449, One-sample t-test: t(486)=26.477, p<0.001) (Fig. 3, Dataset S2). Despite this global pattern, only 65.1% (376/578) of all tested X-chromosomal genes had significantly higher expression from the *Dom* allele. After excluding the 47 genes known to escape X inactivation in normal mouse placenta (Andergassen et al. 2017), 29 genes met our cutoff criteria (see Methods and Dataset S2). Of these, 11 had biallelic expression and 18 had significantly higher expression from the paternal *Spret* allele (Table 3). One *Spret-biased* gene (*Prrg1*) was transgressively overexpressed in hybrid female placenta and *Smarca1*, a gene with biallelic expression on the female X chromosome, was transgressively overexpressed from the maternal (*Dom*) X in hybrid males (Dataset S2).

**Table 3.**
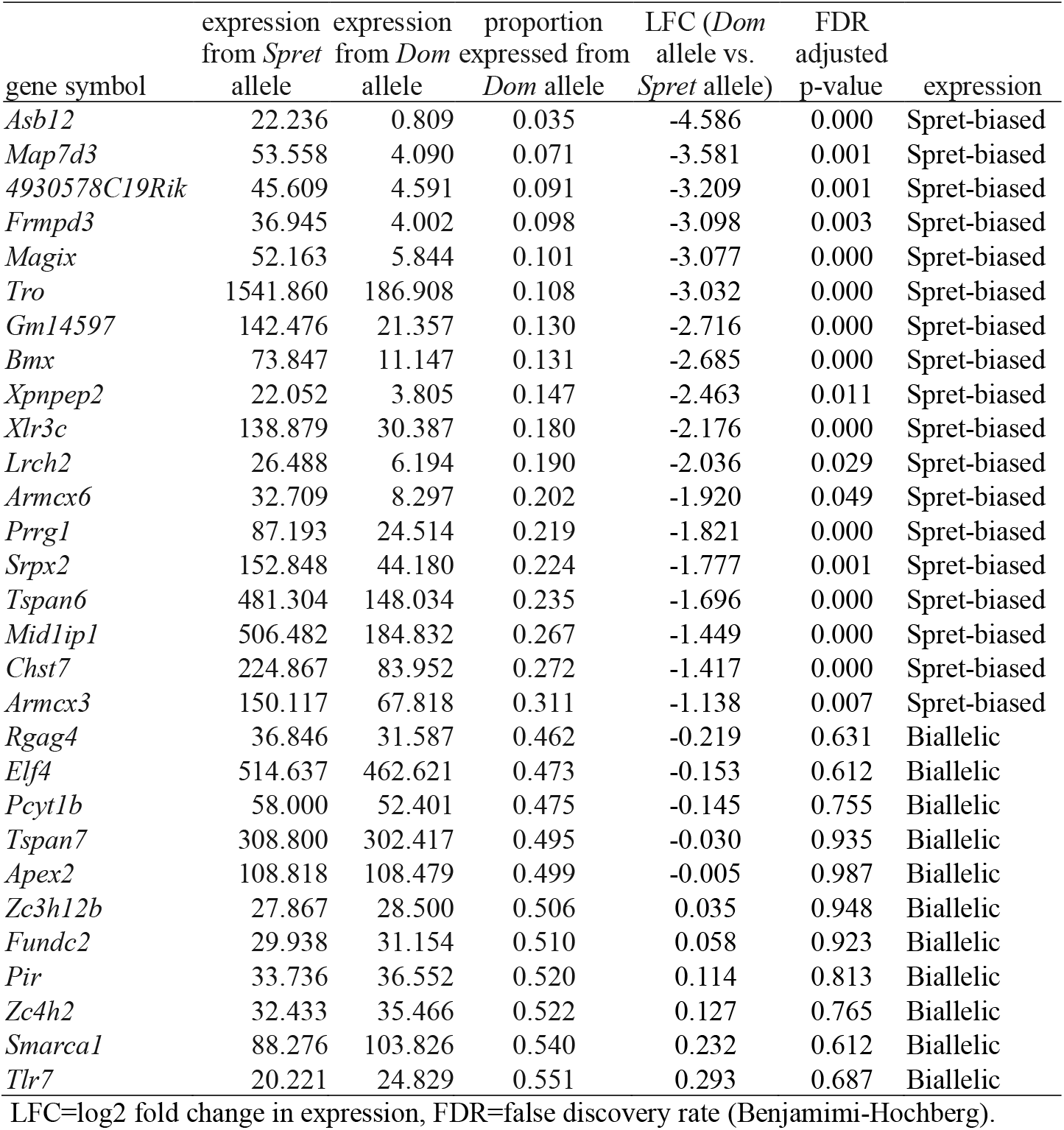
Paternally-biased and biallelic expression on the X chromosome in hybrid females.

| gene symbol | expression<br>from *Spret*<br>allele | expression<br>from *Dom*<br>allele | proportion<br>expressed from<br>*Dom* allele | LFC (*Dom*<br>allele vs.<br>*Spret* allele) | FDR<br>adjusted<br>p-value | expression |
| --- | --- | --- | --- | --- | --- | --- |
| *Asb12* | 22.236 | 0.809 | 0.035 | -4.586 | 0.000 | Spret-biased |
| *Map7d3* | 53.558 | 4.090 | 0.071 | -3.581 | 0.001 | Spret-biased |
| *4930578C19Rik* | 45.609 | 4.591 | 0.091 | -3.209 | 0.001 | Spret-biased |
| *Frmpd3* | 36.945 | 4.002 | 0.098 | -3.098 | 0.003 | Spret-biased |
| *Magix* | 52.163 | 5.844 | 0.101 | -3.077 | 0.000 | Spret-biased |
| *Tro* | 1541.860 | 186.908 | 0.108 | -3.032 | 0.000 | Spret-biased |
| *Gm14597* | 142.476 | 21.357 | 0.130 | -2.716 | 0.000 | Spret-biased |
| *Bmx* | 73.847 | 11.147 | 0.131 | -2.685 | 0.000 | Spret-biased |
| *Xpnpep2* | 22.052 | 3.805 | 0.147 | -2.463 | 0.011 | Spret-biased |
| *Xlr3c* | 138.879 | 30.387 | 0.180 | -2.176 | 0.000 | Spret-biased |
| *Lrch2* | 26.488 | 6.194 | 0.190 | -2.036 | 0.029 | Spret-biased |
| *Armxc6* | 32.709 | 8.297 | 0.202 | -1.920 | 0.049 | Spret-biased |
| *Prrg1* | 87.193 | 24.514 | 0.219 | -1.821 | 0.000 | Spret-biased |
| *Srpx2* | 152.848 | 44.180 | 0.224 | -1.777 | 0.001 | Spret-biased |
| *Tspan6* | 481.304 | 148.034 | 0.235 | -1.696 | 0.000 | Spret-biased |
| *Mid1ip1* | 506.482 | 184.832 | 0.267 | -1.449 | 0.000 | Spret-biased |
| *Chst7* | 224.867 | 83.952 | 0.272 | -1.417 | 0.000 | Spret-biased |
| *Armxc3* | 150.117 | 67.818 | 0.311 | -1.138 | 0.007 | Spret-biased |
| *Rgag4* | 36.846 | 31.587 | 0.462 | -0.219 | 0.631 | Biallelic |
| *Elf4* | 514.637 | 462.621 | 0.473 | -0.153 | 0.612 | Biallelic |
| *Pcyt1b* | 58.000 | 52.401 | 0.475 | -0.145 | 0.755 | Biallelic |
| *Tspan7* | 308.800 | 302.417 | 0.495 | -0.030 | 0.935 | Biallelic |
| *Apex2* | 108.818 | 108.479 | 0.499 | -0.005 | 0.987 | Biallelic |
| *Zc3h12b* | 27.867 | 28.500 | 0.506 | 0.035 | 0.948 | Biallelic |
| *Fundc2* | 29.938 | 31.154 | 0.510 | 0.058 | 0.923 | Biallelic |
| *Pir* | 33.736 | 36.552 | 0.520 | 0.114 | 0.813 | Biallelic |
| *Zc4h2* | 32.433 | 35.466 | 0.522 | 0.127 | 0.765 | Biallelic |
| *Smarca1* | 88.276 | 103.826 | 0.540 | 0.232 | 0.612 | Biallelic |
| *Tlr7* | 20.221 | 24.829 | 0.551 | 0.293 | 0.687 | Biallelic |

LFC=log2 fold change in expression, FDR=false discovery rate (Benjamini-Hochberg).

### Global and X-linked methylation patterns

We assessed genome-wide methylation at CpG islands using reduced representation bisulfite sequencing (RRBS). Throughout this paper, methylation level refers to the mean methylation proportion at CpG islands in promoter regions. In male placentas the autosomal methylation level was 2.38 for *Dom*, 2.37 for hybrids and 2.30 for *Spret*. In female placentas the autosomal methylation level was 2.37 for *Dom*, 2.31 for hybrids and 2.05 for *Spret* (Fig. 4A, Dataset S3). Thus, there was no genome-wide signal of transgressive methylation in hybrids. Consistent with the generally repressive role of methylation in promoter regions, there was a weak negative correlation between global methylation and expression for all datasets (Females: Hybrid: R^2^adj=0.115, p<0.001, *Dom:* R^2^adj=0.118, p<0.001 *Spret:* R^2^adj=0.107, p<0.001; Males: Hybrid: R^2^adj=0.128, p<0.001, *Dom*: R^2^adj=0.127, p<0.001, *Spret*: R^2^adj=0.12, p<0.001) (Fig. S4). Mean methylation levels on the X chromosome were 6.19 for *Dom* males, 7.23 for *Spret* males and 6.99 for hybrid males. In hybrid male vs. *Dom* male placentas, the mean LFC in methylation levels on the *M. m. domesticus* X was significantly higher than the mean LFC in autosomal methylation levels (mean LFC_X_=0.36, mean LFC_Autosomes_=0.10; One-sample t-test: t(172)= −2.17, p= 0.03). Of 170 X-linked genes assessed, 17.05% (29/170) were significantly hypermethylated in hybrid vs. *Dom* males whereas 6.47% (11/170) were hypomethylated. Compared to *Spret* males, 9.41% (16/170) of X-linked genes were hypermethylated and 14.11% (24/170) were hypomethylated in hybrid male placenta.

**Figure 4.**
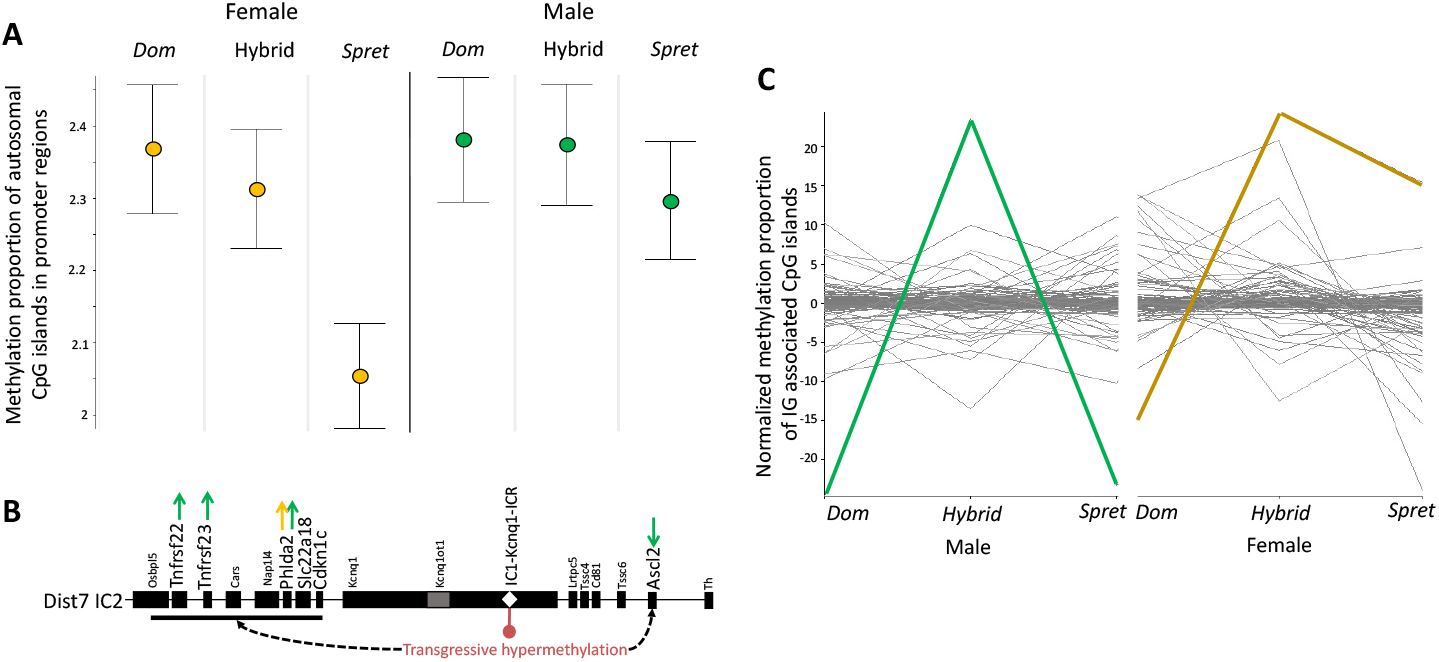
Methylation patterns of CpG islands in gene promoters and the *Kcn1q* imprinting control region (ICR). Male data depicted in green and female data in yellow throughout the figure. (A) The mean proportions of methylated reads for CpG islands in autosomal promoter regions. (B) Schematic representation of the distal chromosome 7 imprinting cluster 2 (Dist7-IC2) with the ICR within the gene *Kcn1q* marked with a white diamond. Genes with transgressive expression are in larger font; arrows indicate the direction of expression in hybrid males (green) and females (gold). (C) Line plots show the normalized proportions of methylated reads for all IG promoter or ICR-associated CpG islands, where each line represents an IG or ICR-associated CpG island. The IC2-Kcnq1-ICR CpG island is marked in green in the male plot and gold in the female plot. Error bars in (A), standard error.

We also assessed allele specific methylation (ASM) for male and female hybrids. In hybrid male placentas the autosomal methylation level was 2.18 for the *Dom*-allele and 2.15 for the *Spret*-allele. In hybrid female placentas the autosomal methylation level was 2.16 for the *Dom*-allele and 2.13 for the *Spret*-allele. As expected, the methylation level of the maternal X chromosome in female hybrids was lower compared to the paternal X (X_dom_=12.03, X_spret_=28.92) (Dataset S3).

### Differential methylation of gene promoter CpGs in the hybrid placenta

For the analysis of differential methylation, a ≥ 1.5 fold difference in methylation proportion (≥ 0.5 LFC) with an FDR value ≤ 0.05 was considered significant. In hybrid male placenta, 2.59% (176/6,794) of all tested genes’ promoter regions were hypermethylated and 2.05% (139/6,794) were hypomethylated compared to *Dom* male placenta; 2.87% (195/6,794) were hypermethylated and 2.05% (139/6,795) were hypomethylated compared to *Spret* male placenta (Dataset S3). In hybrid female placenta, 1.46% (99/6,798) of all tested genes’ promoter regions were hypermethylated and 1.43% (97/6,798) hypomethylated compared to *Dom* female placenta; 2.69% (183/6,798) were hypermethylated and 1.35% (92/6,798) were hypomethylated compared to *Spret* female placenta (Dataset S3). In hybrid male placenta, 23 genes were transgressively hyper-, and 6 transgressively hypomethylated. In female hybrids, 15 genes were transgressively hypermethylated and 1 was hypomethylated. There was no overlap in transgressive promoter methylation between the sexes (Table S1).

### Transgressive methylation of imprinted gene-associated CpGs in the hybrid placenta

We tested for transgressive methylation in CpG islands associated with IGs using the same criteria for significance as for genome-wide differential methylation. In hybrid male placentas, 4 IG-associated CpG islands were transgressively methylated: *Igf2r, Phlda2, Trmp5*, and *Kcnq1*. The latter 3 are located in the dist7 imprinting cluster and include hypermethylation of the CpG island in *Kcnq1* intron 10, the cluster’s imprinting control region (ICR2) (Lee et al. 1999) (Fig. 4B, Table 4). The CpG associated with this ICR, which is normally maternally methylated (Zhang et al. 2014), was strongly hypermethylated in hybrid males and, to a lesser extent, in hybrid females compared to all other IG and ICR-associated CpGs for which methylation was measured (Fig. 4C). A *Peg12*-associated CpG island was transgressively hypermethylated in hybrid female placentas (Table 4).

**Table 4.** Transgressive imprinted gene-associated CpG island methylation in male and female hybrids.

|  | Chr. | closest IG | % reads methylated in CpG island |  |  | LFC Hybrid<br>vs. *Dom* | LFC Hybrid<br>vs. *Spret* | FDR Hybrid<br>vs. *Dom* | FDR Hybrid<br>vs. *Spret* |
| --- | --- | --- | --- | --- | --- | --- | --- | --- | --- |
|  |  |  | *Dom* | Hybrid | *Spret* |  |  |  |  |
| male<br>placenta | **17** | ***Igf2r*** | **40.33** | **27.35** | **45.07** | **-0.56** | **-0.72** | **0.015** | **0.001** |
|  | **7** | ***Phlda2*** | **5.68** | **3.22** | **5.41** | **-0.82** | **-0.75** | **<0.001** | **0.011** |
|  | **7** | ***Trpm5*** | **18.07** | **10.09** | **24.28** | **-0.84** | **-1.27** | **<0.001** | **<0.001** |
|  | **7** | ***Kcnq1*** | **21.40** | **69.62** | **22.78** | **1.70** | **1.61** | **<0.001** | **<0.001** |
| female<br>placenta | 6 | *Peg10* | 52.79 | 28.23 | 38.15 | -0.90 | -0.43 | <0.001 | 0.001 |
|  | 12 | *Begain* | 48.25 | 33.72 | 42.68 | -0.52 | -0.34 | <0.001 | <0.001 |
|  | **7** | ***Peg12*** | **5.70** | **8.91** | **4.78** | **0.64** | **0.90** | **<0.001** | **<0.001** |
|  | 17 | *Igf2r* | 53.46 | 36.33 | 42.35 | -0.56 | -0.22 | <0.001 | <0.001 |
|  | 7 | *Kcnq1* | 30.98 | 70.44 | 61.44 | 1.18 | 0.20 | 0.017 | 0.002 |

Chr.=Chromosome, IG=imprinted gene, LFC=log 2 fold change in expression. Transgressively methylated CpG islands in bold, FDR=false discovery rate (Benjamini-Hochberg)

### Allele specific methylation

We evaluated allele specific methylation (ASM) in hybrid placentas by testing for allelic differences in methylation on promoter region CpGs. We considered log2 fold change (LFC) in methylation ≥ 1 (2 times higher or lower methylation from one allele compared to the other), and Benjamini-Hochberg-corrected p ≤ 0.05 as significant ASM. In hybrid male placentas, 1.85% and 0.9% of all tested autosomal promoter regions had significantly higher methylation levels on the *Dom* or *Spret* allele, respectively (*Dom* ASM: 68/3,668, including IGs *Slc22a3, Cdnk1c* and *Ascl2; Spret* ASM: 33/3,668, including IG *Gnas*) (Dataset S3). In hybrid female placentas, 1.15% and 0.55% of all tested autosomal promoter regions had significantly higher methylation levels on the *Dom* or *Spret* allele, respectively (*Dom* ASM: 42/3,668, including IGs *Trpm5, Slc22a3, Ascl2* and *Phlda2; Spret* ASM: 20/3,668, including IG *Gnas*) (Dataset S3).

### X-linked and imprinted gene network analyses

We used gene network analysis to test for functional associations between transgressively expressed genes and 1) X-linked genes with *Spret*-biased or biallelic expression in hybrid female placenta (*n*=29), and 2) placental imprinted genes that retained allele specific expression in hybrids (*n*=33). The 665 interactor genes for the X-linked gene list were not enriched for genes with transgressive expression in male or female hybrids (Fisher’s exact test: Males: p=0.492, odds ratio = 0.766, Females: p=1, odds ratio = 0.903); 11 and 4 were transgressively expressed in hybrid male and female placentas, respectively (Dataset S2). Similarly, IG interactors were not significantly enriched for transgressively expressed genes in either sex (Fisher’s exact test: Males: p=0.277, odds ratio = 0.542, Females: p=0.412, odds ratio = 1.3). Of the 553 IG interactors, 6 and 4 were uniquely transgressively expressed in hybrid male and female placentas, respectively, and 4 were transgressive in both sexes (Dataset S2).

## DISCUSSION

Imprinted genes are critical regulators of prenatal growth and development in mammals (Ferguson-Smith 2011). This function, together with characteristic features of regulation and expression, makes imprinted genes compelling candidates for asymmetric growth abnormalities in mammalian hybrids. We used transcriptome and reduced representation bisulfite sequencing to evaluate the contribution of IGs to a long-standing example of parent-of-origin placental growth dysplasia in hybrid mice. We found 1) transgressive expression and DNA methylation in an imprinted gene cluster that is a strong candidate locus for transgressive placental growth, 2) more extreme undergrowth and more transgressive expression and methylation in male vs. female hybrid placentas, and 3) an excess of genes that escape imprinted X chromosome inactivation in hybrid females. We discuss these results in light of prior work on this and other cases of hybrid growth dysplasia, and address the perplexing question of why hybrid placental phenotypes are more extreme in the heterogametic sex.

### Placental defects and inviability in hybrid mice

The first description of hybrid developmental abnormalities in the cross between female *M. m. domesticus* and male *M. spretus* (*Dom* x *Spret*) reported placental undergrowth relative to *M. m. domesticus* (Zechner et al. 1996) but did not make comparisons to the smaller parental species, *M. spretus*. Here, we show that hybrid placental weight is significantly reduced relative to both parental species, with more extreme undergrowth in males. Placental histology is also consistent with the original report, in which the spongiotrophoblast layer (the placenta’s endocrine compartment) was reduced in hybrid relative to *M. m. domesticus* placentas (Zechner et al. 1996). We find that the junctional zone, of which the spongiotrophoblast layer is the main component (Coan et al. 2006; John 2013), is qualitatively narrower in hybrid placenta relative to both parental species. Notably, we find a moderate but statistically significant reduction in the viability of hybrid conceptuses, providing the first quantitative evidence that developmental abnormalities contribute to postzygotic reproductive isolation in this direction of the cross.

### The contribution of imprinted genes to hybrid placental growth dysplasia

Prior analysis of allelic expression and DNA methylation for 18 candidate IGs in various adult F1 hybrid tissues found patterns suggestive of partial loss of imprinting (biallelic expression) in both directions of the *M. m. domesticus/M. spretus* cross (Shi et al. 2005). In the placenta, however, tests for the contribution of two paternally expressed IGs to hybrid growth dysplasia were inconclusive (Zechner et al. 2002; 2004). Excess Igf2 protein was observed in overgrown placentas but the overgrowth phenotype persisted in the absence of functional Igf2 (Zechner et al. 2002); *Peg3* retained normal expression from the paternal allele in hybrid placenta and the locus was not associated with placental abnormalities in a backcross mapping panel (Zechner et al. 2004).

Using whole transcriptome data, we show that the typical parent-of-origin expression patterns of most placental imprinted genes are retained in *Dom* x *Spret* F1 hybrids. Moreover, the expression of three IGs with evidence of biallelic expression in hybrid placenta is within the range of both parental species. Thus, loss of imprinting and associated altered dosage of imprinted genes cannot explain transgressive placental undergrowth in *Dom* x *Spret* hybrids. Similarly, we did not find widespread transgressive expression of imprinted genes, nor did functional associations with canonically expressed (i.e., monoallelic) imprinted genes explain a significant proportion of all transgressive expression. Instead, transgressively expressed IGs were concentrated in the *Kcnq1* cluster on distal chromosome 7 in association with hypermethylation of the cluster’s imprinting control region, ICR2. In humans, loss of methylation on ICR2, and consequent silencing of the maternally expressed genes in the *KCNQ1* cluster, is a common cause of Beckwith-Wiedemann syndrome, an imprinting disorder characterized by fetal and placental overgrowth (Lee et al. 1999; Cooper et al. 2017). This association between ICR2 hypomethylation and conceptus overgrowth implicates ICR2 hypermethylation as an effector of the placental undergrowth phenotype studied here. Indeed, functional studies in lab mice indicate that misexpression of just two maternally expressed genes in the *Kcnq1* cluster, *Phlda2* (overexpressed in hybrids of both sexes) and *Ascl2* (underexpressed in males), could underlie placental undergrowth in hybrids, and the more extreme male phenotype.

Overexpression of *Phlda2* in transgenic mice results in both fetal and placental growth restriction, with placentas characterized by a reduced junctional zone and deficits in trophoblast glycogen cells (Tunster et al. 2016a), a putative fetal energy source in late gestation (Coan et al. 2006). Whereas *Ascl2* (formerly *Mash2*) knockouts die in midgestation (e10) due to complete failure of spongiotrophoblast layer formation (Guillemot et al. 1994; Tanaka et al. 1999),underexpression of *Ascl2* causes non-lethal placental undergrowth characterized by reduced spongiotrophoblast and a lack of trophoblast glycogen cells together with an expanded giant cell layer and disorganization of vasculature critical to feto-maternal exchange (Oh-McGinnis et al. 2011). This placental phenotype bears histological resemblance to that in *Dom* x *Spret* hybrids (Fig S1; Zechner et al. 1996). Notably, reduction in *Ascl2* expression in the transgenic model also causes upregulation of *Phlda2*, suggesting an upstream regulatory effect of *Ascl2* on *Phlda2* (Oh-McGinnis et al. 2011). Collectively, these and other functional studies of *Phlda2* and *Ascl2* in mouse indicate that the essential role of *Ascl2* in placental development includes modulation of the suppressive effects of *Phlda2* on spongiotrophoblast expansion (Tunster et al. 2016b). Application of this model to placental undergrowth in hybrids suggests that moderate *Phlda2* overexpression is sufficient to cause observed placental growth restriction in females, whereas the addition of *Ascl2* underexpression in males increases the severity of the growth phenotype in association with more extreme upregulation of *Phlda2*. Given the availability of transgenic mouse models for both *Ascl2* and *Phlda2* (e.g. Tunster et al. 2016a,b; Creeth et al. 2018), it may be possible to test the proposed effects of altered dosage on hybrid placenta using crosses between transgenic females and unmanipulated *M. spretus* males.

Interestingly, in the cross between female *Peromyscus polionotus* and male *P. maniculatus*, *Kcnq1* is one of three major IG clusters that are dysregulated in overgrown placentas (Duselis and Vrana 2007; Wiley et al. 2008). In particular, *Phlda2* is downregulated (Duselis and Vrana 2007) in association with loss of methylation on ICR2 (Wiley et al. 2008). Given that opposite patterns of altered methylation at this locus are associated with opposite placental growth dysplasia phenotypes in hybrids from distantly related rodent genera, we predict that hypomethylation of *Kcnq1* ICR2 will emerge as a major determinant of hybrid placental overgrowth in the reciprocal *Spret* x *Dom* cross. More generally, the association between altered methylation on *Kcnq1* ICR2 and prenatal growth dysplasia in both rodent hybrids and humans suggests that, whether due to intrinsic incompatibilities in hybrids or *de novo* epimutations, the locus is particularly vulnerable to epigenetic perturbation.

### Haldane’s rule and the large X effect in hybrid placenta

In agreement with prior studies of the *M. m. domesticus/M. spretus* cross (e.g. Zechner et al. 1996; Hemberger et al. 2001), we found that hybrid placental phenotypes obey Haldane’s rule: *Dom* x *Spret* hybrid male placentas are significantly smaller than those of their already growth restricted sisters. Moreover, there were more than twice as many transgressively expressed genes in hybrid males and, even within the *Kcnq1* cluster, transgressive expression and methylation were greater in hybrid males. Importantly, while male-limited underexpression of *Ascl2* may be the direct cause of the more extreme male growth phenotype, and may account for some of the excess transgressive expression in males due to imbalanced proportions of *Ascl2*-regulated placental cell lineages, sex-specific dysregulation of autosomal genes requires additional explanation.

Of the genetic hypotheses for Haldane’s rule, all that apply to hybrid inviability involve large negative effects of X- (or Z-) linked incompatibilities in the heterogametic sex, with the predominant explanation being that most DMI loci act as recessives or partial recessives in hybrids and are therefore uniquely exposed on the X chromosome in the heterogametic sex (Turelli and Orr 1995; Orr and Turelli 1996; reviewed in Coyne and Orr 2004; Delph and Demuth 2016). This “large X effect” (Coyne and Orr 1989) is further increased if DMIs accumulate preferentially on the X chromosome relative to the autosomes (Charlesworth et al. 1987), as is the case for loci involved in hybrid male sterility in both *Drosophila* and mice (Masly and Presgraves 2007; Good et al. 2008; Larson et al. 2017).

Consistent with a large X effect on hybrid placental phenotypes in the *M. m. domesticus*/*M. spretus* cross, backcross mapping studies found a large contribution of the X chromosome to placental growth dysplasia, with undergrowth associated with *M. m. domesticus* genotypes and overgrowth with *M. spretus* genotypes (Zechner et al. 1996; Hemberger et al. 1999, 2001). However, because imprinted XCI in normal (i.e. non-hybrid) rodent placenta results in functional hemizygosity for the maternal X chromosome in both sexes, adherence to Haldane’s rule in this tissue is unexpected. We consider four potential explanations: XCI escape in females, X-Y interactions in males, *trans*-acting effects of altered chromatin structure on the X chromosome in males, and sexually dimorphic maternal effects.

First, partial dysregulation of imprinted XCI, resulting in biallelic expression of some X-linked genes, could buffer females from the effects of X-linked incompatibilities that are fully exposed in males. This explanation for hybrid male-biased placental growth dysplasia was proposed in the original description of the phenotype (Zechner et al. 1996) but later dismissed when three X-linked markers showed the normal pattern of maternal expression/paternal silencing in hybrid female placenta (Hemberger et al. 2001). We found that, while allelic expression for the majority of X-linked genes retains strong maternal bias, 29 genes that are not known to escape XCI in mouse placenta met our criteria for having biallelic or paternally-biased expression in hybrid female placenta. None of these genes are obvious candidates for direct effects on placental growth and development, and genes with transgressive expression in male placenta were not globally enriched among functional interactors for the 29 X-linked genes. However, among the genes with biallelic expression in females, *Smarca1* is a candidate for regulatory effects on hybrid males. *Smarca1* encodes a chromatin remodeling enzyme that suppresses cell proliferation and attenuates Wnt-signaling (Eckey et al. 2012), processes crucial for embryogenesis and placental development (Parr et al. 2001; Logan and Nusse 2004). Members of the Smarca gene family promote insulator protein CTCF binding with downstream effects on CTCF-regulated genes (Wiechens et al. 2016), which include several imprinted gene clusters (Lléres et al. 2019). Given that *Smarca1* was transgressively overexpressed in hybrid male placenta, the potential contribution of this gene to the difference between hybrid male and female phenotypes is worthy of further study.

Second, the opportunity for negative epistatic interactions between the X and Y chromosomes is unique to hybrid males. In the absence of support for alternative explanations, Hemberger and colleagues proposed that X-Y interactions account for adherence to Haldane’s rule in the *M. m. domesticus*/*M. spretus* cross (Hemberger et al. 2001). The current data do not address this hypothesis; a test would require generation of Y-introgression lines for crosses in which X and Y chromosomes derive from the same parent species in hybrid males. However, we think it is unlikely to explain male-biased placental deficits. Whereas negative X-Y interactions are important in a handful of cases of Haldane’s rule for hybrid male sterility (Coyne et al. 2004; Mishra and Singh 2007) there are, to our knowledge, no published examples of this genetic architecture underlying preferential inviability of the heterogametic sex in animals. In *Dom* x *Spret* hybrids, transgressive placental phenotypes are in the same direction in both sexes, albeit more extreme in males, and X chromosome genotype has a large effect on growth dysplasia in both sexes (Zechner et al. 1996; Hemberger et al. 1999, 2001). Therefore, a male-specific genetic architecture for which there is minimal precedent in other systems does not provide a parsimonious explanation for the observed data.

Third, sex differences in chromatin structure on the X could have *trans*-acting effects on autosomal expression that differ between male and female hybrids (Hemberger et al. 1999). Sex chromosome complement explains sex differences in autosomal gene expression that are independent of gonadal sex and associated hormonal differences between mammalian males and females (Bermejo-Alvarez et al. 2010; Raznahan et al. 2018). Moreover, the parental origin of the single X chromosome in humans with Turner syndrome (45,X) impacts prenatal survival and the severity of postnatal phenotypes, indicating parent-of-origin epigenetic effects (Sagi et al. 2007; Grande et al. 2019). Circumstantial evidence supporting the hypothesis that aberrant epigenetic effects of the X chromosome contribute to male-biased placental growth dysplasia includes our finding of elevated mean LFC in methylation levels on the *M. m. domesticus* X chromosome in hybrid vs. *Dom* males and the fact that, in crosses using subcongenic mice with intervals of the *M. spretus* X on an *M. m. domesticus* autosomal background, the degree of placental overgrowth increases with the size, rather than the location, of the *M. spretus*-derived interval (Hemberger et al. 1999).

Fourth, we would expect male-biased sensitivity to any negative effect of a hybrid pregnancy on the maternal intrauterine environment. Maternal stress during pregnancy, whether physiological or psychological, negatively impacts both sexes (Davis and Pfaff 2014). However, the immediate consequences of an adverse intrauterine environment are generally more severe in males, with more pronounced placental pathology and intrauterine growth restriction, and a higher rate of mortality for male fetuses (Cooperstock and Campbell 1996; Walker et al. 2012; Sandman et al. 2013; Davis and Pfaff 2014). Female *M. m. domesticus* carrying hybrid litters are potentially exposed to altered placental endocrine signaling due to deficits in spongiotrophoblast cells in hybrid placentas (Zechner et al. 1996), and exhibit altered neural gene expression at e17.5 (Arévalo and Campbell 2020) and reduced maternal behavior immediately postpartum (Gardner et al. 2019). These observations suggest that hybrid pregnancy disrupts maternal homeostasis and, in this sense, is a physiological stressor. Moreover, maternal immunotolerance of the conceptus is mediated by the placenta and maternal immune response to male fetuses may be greater than that to female fetuses (Kahn and Baltimore 2010; Bogaert et al. 2018). Maternal immune response to males could be elevated by heterospecific Y-linked gene expression in placenta in hybrid pregnancies.

We emphasize that none of the mechanisms proposed here to explain Haldane’s rule in hybrid placenta are mutually exclusive. For example, intrinsic negative effects of hybrid male sex chromosome complement on autosomal gene expression, whether epigenetic or genetic, could be amplified by male-biased sensitivity to an adverse maternal environment and/or reduced maternal immunotolerance of hybrid male conceptuses. Finally, it is noteworthy that 18 X-linked genes have effectively hemizygous expression from the paternal (*M. spretus*) allele in hybrid female placenta. Given the large contribution of the *M. spretus* X chromosome to overgrowth phenotypes in the reciprocal cross (Zechner et al. 1996; Hemberger et al. 1999, 2001) it is possible that the apparent reduction in the severity of placental undergrowth in hybrid females relative to males actually reflects the opposing effects of incompatibilities involving the *M. spretus* X chromosome, to which males are not exposed.

### Conclusions

The results presented here provide new insight into the genetic basis of hybrid developmental abnormalities and add to prior evidence that placental imprinted genes play a key role in growth dysplasia and associated inviability in mammalian hybrids (Vrana 2000; Brekke and Good 2016). In particular, we find hypermethylation of the *Kcnq1* ICR2 in hybrid placenta, in conjunction with transgressive expression of several maternally expressed IGs in this cluster. Importantly, the direction of *Phlda2* and *Ascl2* misexpression, and of placental growth phenotypes, parallels that in transgenic mouse models for the same genes. This suggests that epigenetic dysregulation of few imprinted genes can have large negative effects on hybrid placental development.

Given that there is also a large X effect on placental abnormalities in the *M. m. domesticus*/*M. spretus* cross (Zechner et al. 1996; Hemberger et al. 1999), it is tempting to infer a simple genetic architecture in which IGs are the autosomal interaction partners in X-linked incompatibilities. Assuming asymmetric fitness optima for maternally and paternally inherited alleles that influence conceptus growth, Patten (2018) proposed that imprinted XCI favors antagonistic regulatory effects of the maternally inherited X chromosome on paternally expressed IGs in placenta. If this parental conflict is recurrent and lineage-specific, within-species equilibrium could be disrupted when parents belong to different species. Negative epistasis between the maternal X and at least one paternally expressed IG in overgrown *Peromyscus* hybrids (Vrana et al. 1998, 2000; Losciavo et al. 2007) and widespread underexpression of Pegs in overgrown *Phodopus* hybrids (Brekke et al. 2016) provide some support for this proposition. However, incompatibilities involving the maternally inherited *M. m. domesticus* X chromosome and the maternally expressed imprinted genes highlighted here are implausible. Moreover, transgressive expression of *Phlda2* and *Ascl2* is likely a direct consequence of excess methylation on the cluster’s imprinting control region. Whether this local epigenetic perturbation is a consequence of X-linked or other genic incompatibilities, or a byproduct of epigenetic instability in hybrid genomes, is currently unclear. Genome-wide QTL and eQTL mapping will be a critical first step in reconciling the large role of the X, adherence to Haldane’s rule, and the contribution of IGs to hybrid placental growth dysplasia in the *M. m. domesticus*/*M. spretus* cross.

## Acknowledgements

We thank B. Horn and S. Windle for assistance with mouse husbandry and J. Good for the gift of the SFM/Pas strain. This work was supported by the National Science Foundation (NSF-IOS 1558109 to P.C).

