## Supplemental Figures and Tables for "Haldane’s rule in the placenta: sex-biased misregulation of the *Kcnq1* imprinting cluster in hybrid mice"

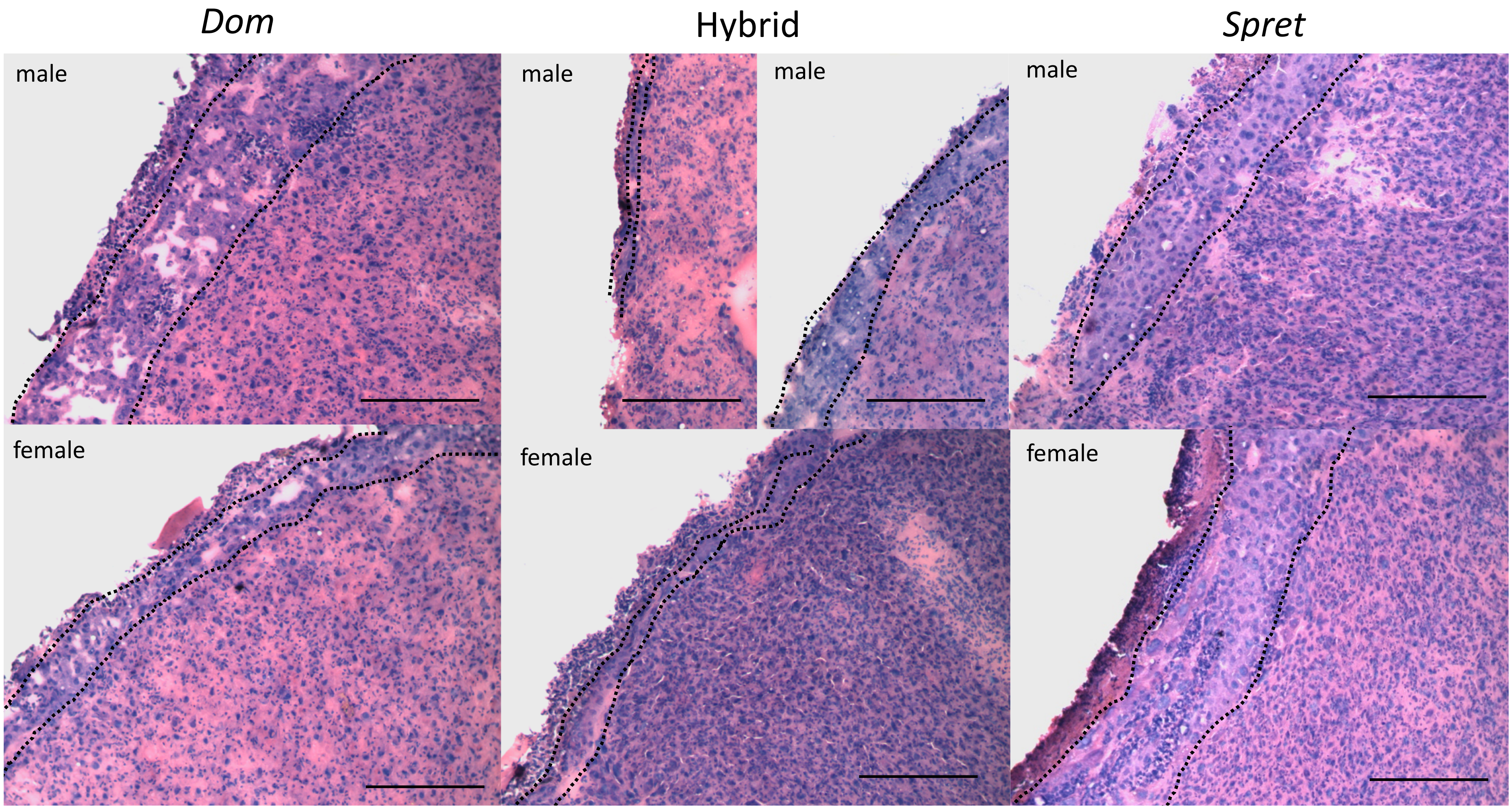


Figure S1. Representative images of histological placenta sections. Cryosections at 20 microns stained with H&E, scale bars represent 400µm. Approximate borders of the junctional zone are marked in each image. *Dom = Mus musculus domesticus*, *Spret = Mus spretus*. The decidual layer was removed (see Methods).

**A**

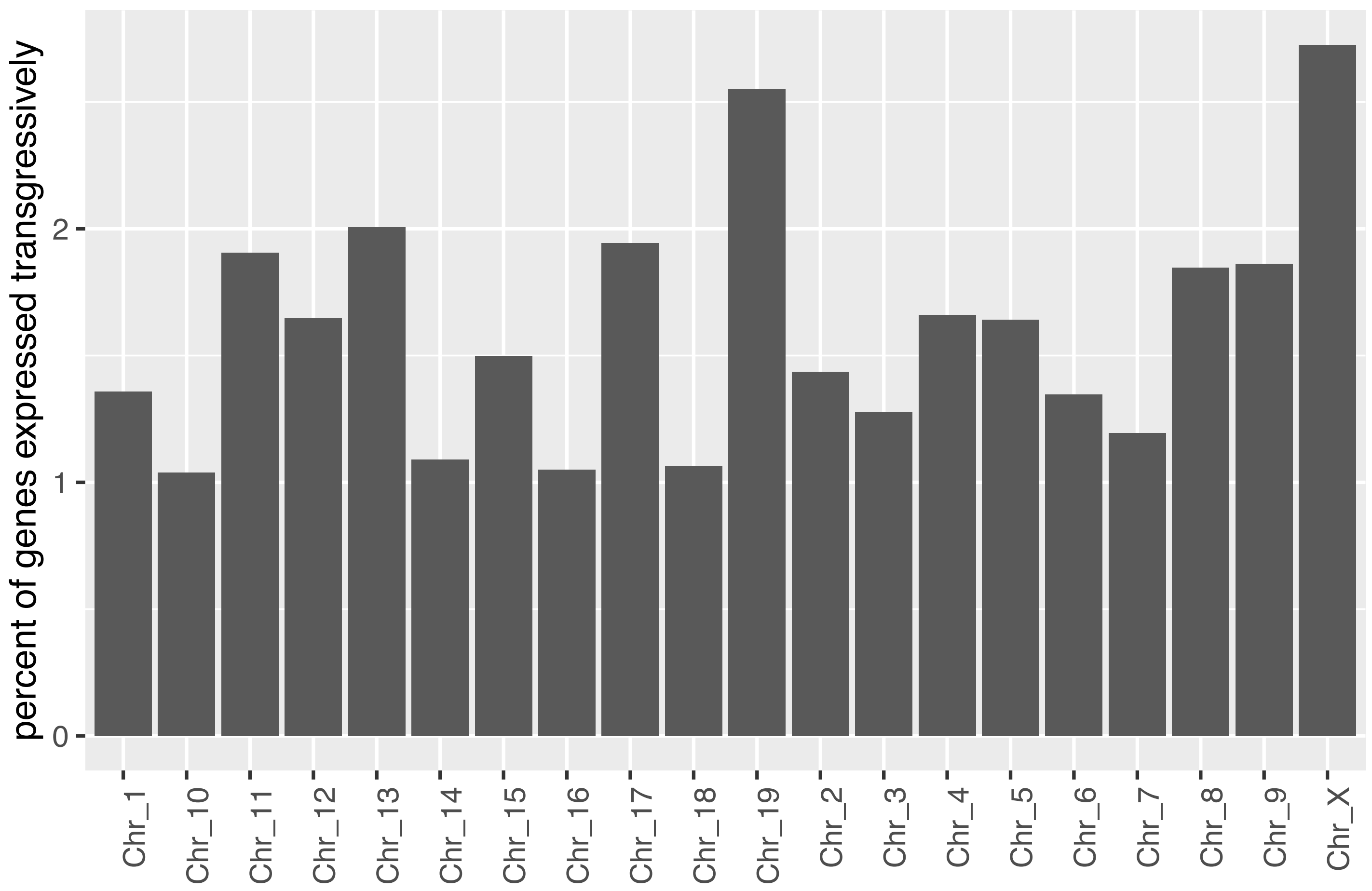


**B**

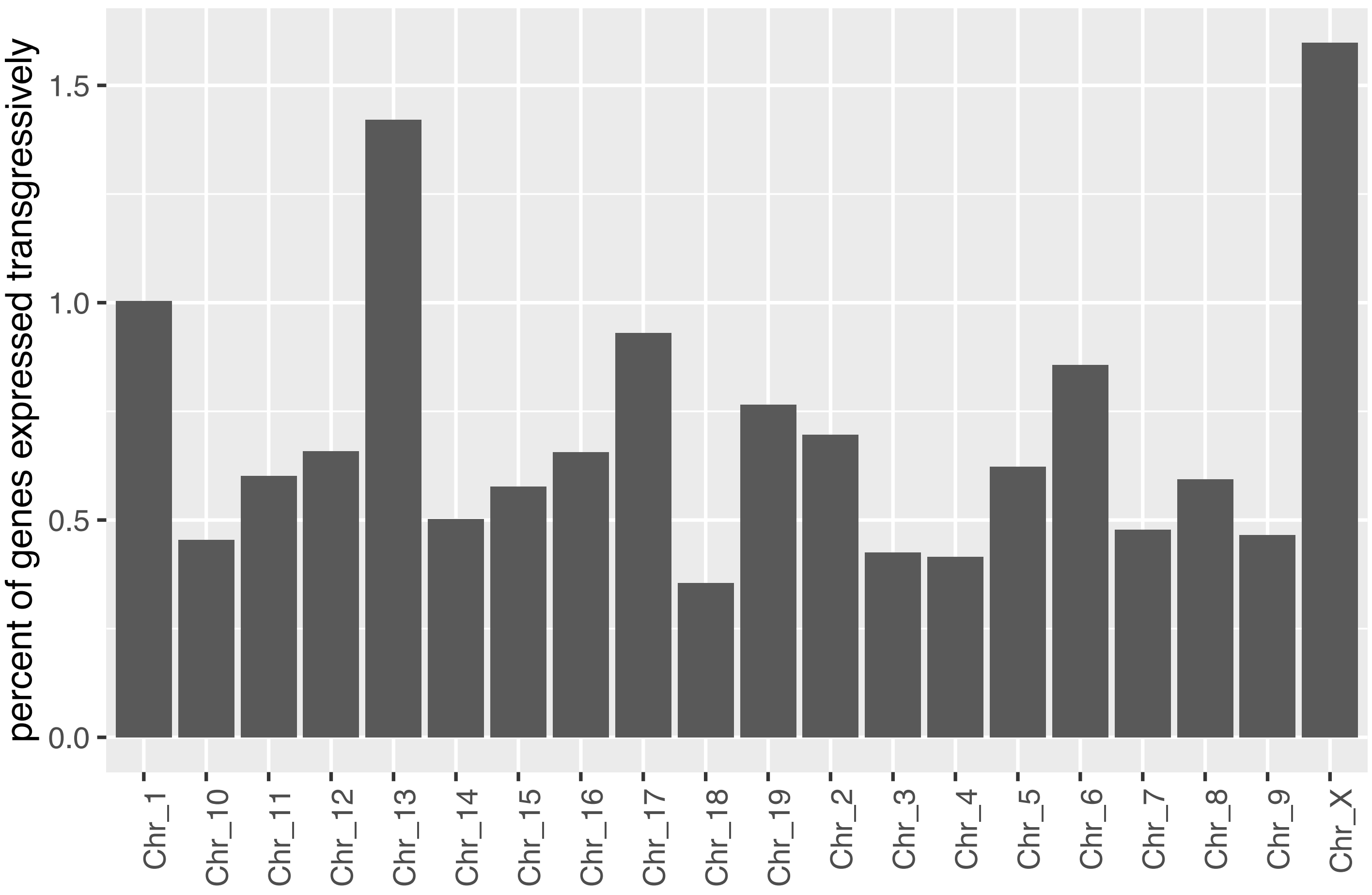


Figure S2. Barplots showing the percentage of genes expressed transgressively on each chromosome for (A) males and (B) females. Only chromosomes that contain transgressively expressed genes are shown.

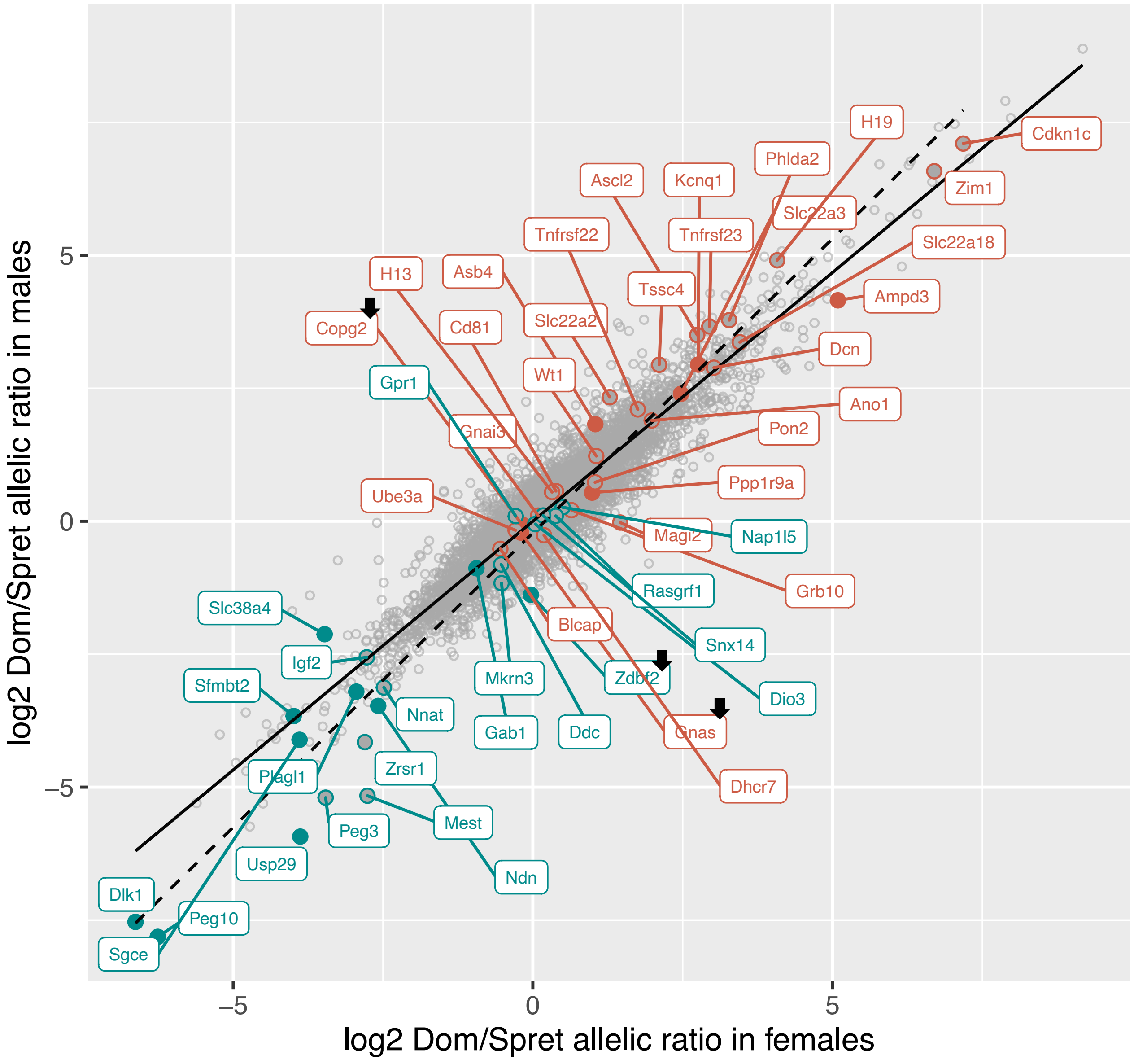


Figure S3. Scatterplot showing the relationship between male and female allelic expression ratios with maternally expressed imprinted genes in red and paternally expressed imprinted genes in blue. Genes with previously validated imprinted expression in e17.5 placenta (Babak et al. 2015) are indicated with filled circles. Imprinted genes with loss of imprinting in hybrids are indicated with an arrow.

**A**

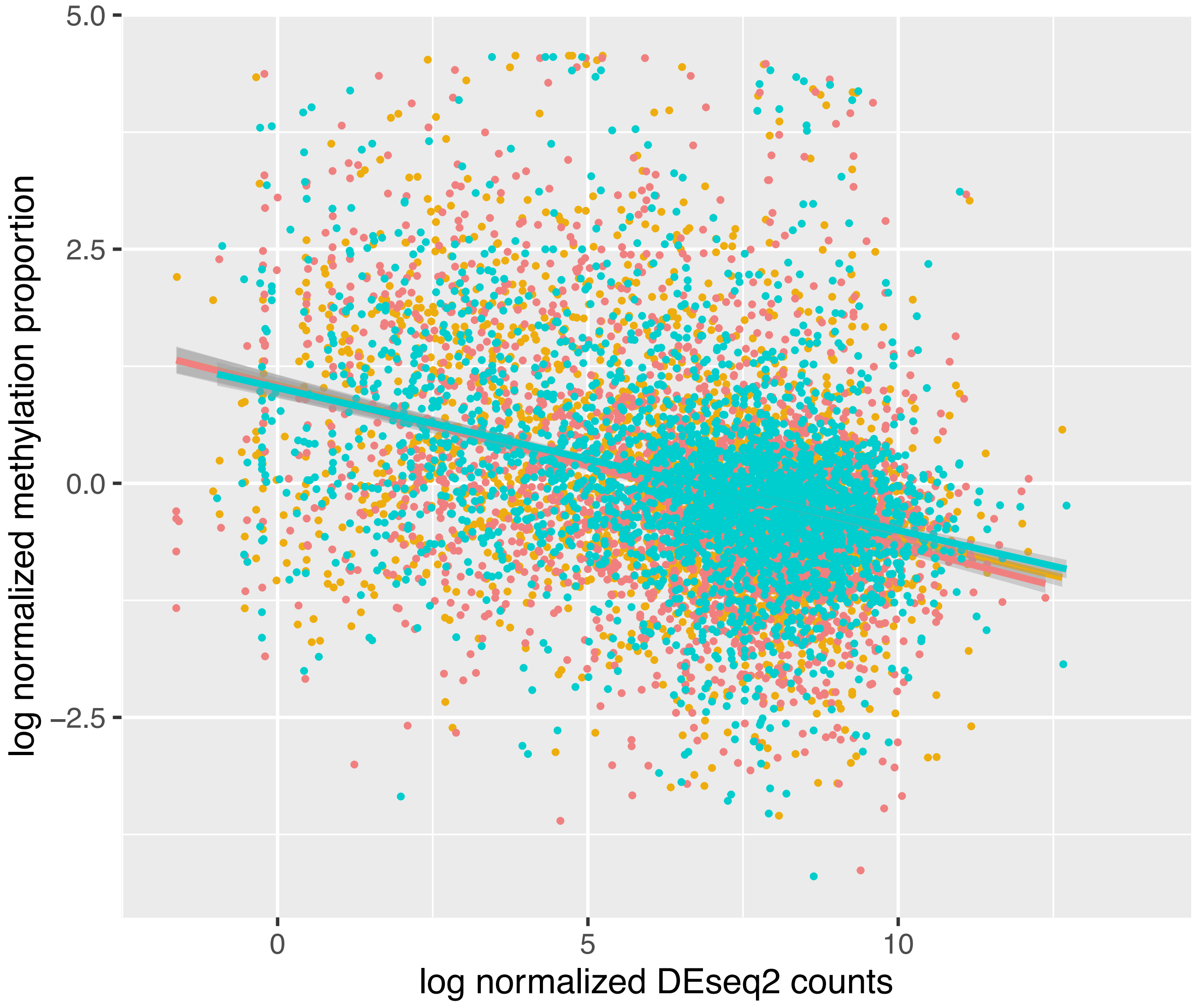


**B**

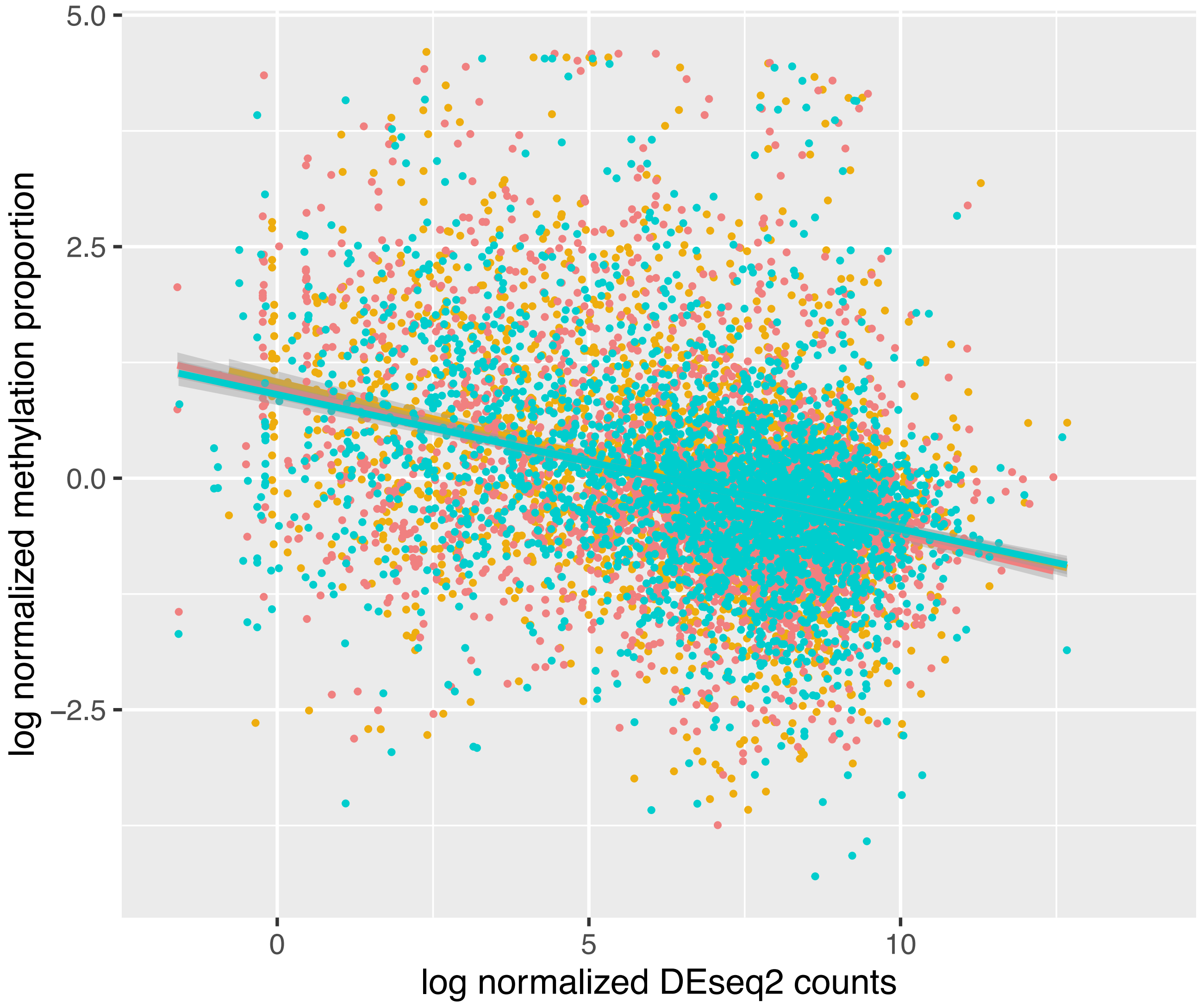


Figure S4: Scatterplots showing the correlation between gene promotor CpG methylation and gene expression for (A) males and (B) females. *Dom,* red; Hybrid, gold; *Spret,* cyan.

Table S1. Transgressive CpG island methylation located upstream of genes in hybrids

| Genotype | Chr. | Downstream Gene | Ensembl ID | Percentage of reads methylatd in CpG island *Dom* | *Hybrid* | *Spret* | LFC Hybrid vs. *Dom* | LFC Hybrid vs. *Spret* | FDR Hybrid vs. *Dom* | FDR Hybrid vs. *Spret* |
|---|---|---|---|---|---|---|---|---|---|---|
| | 16 | Ccdc116 | ENSMUSG00000022768 | 2.64 | 0.98 | 3.99 | -1.43 | -2.02 | 0.04 | 0.00 |
| | 10 | Irak3 | ENSMUSG00000020227 | 2.04 | 0.83 | 3.33 | -1.30 | -2.01 | 0.00 | 0.00 |
| | 4 | Ski | ENSMUSG00000029050 | 1.99 | 0.91 | 2.52 | -1.12 | -1.46 | 0.01 | 0.04 |
| | 4 | Tnfrsf25 | ENSMUSG00000024793 | 37.62 | 20.03 | 40.13 | -0.91 | -1.00 | 0.00 | 0.00 |
| | 1 | Bcl2 | ENSMUSG00000057329 | 27.96 | 18.15 | 27.42 | -0.62 | -0.59 | 0.02 | 0.00 |
| | 11 | Slc13a5 | ENSMUSG00000020805 | 9.21 | 6.31 | 9.14 | -0.55 | -0.53 | 0.02 | 0.03 |
| | 6 | Nxph1 | ENSMUSG00000046178 | 8.84 | 12.65 | 6.99 | 0.52 | 0.86 | 0.00 | 0.00 |
| | 9 | Pde4a | ENSMUSG00000032177 | 36.91 | 53.65 | 37.09 | 0.54 | 0.53 | 0.01 | 0.04 |
| | 11 | Tlx3 | ENSMUSG00000040610 | 4.44 | 7.14 | 5.03 | 0.68 | 0.51 | 0.00 | 0.01 |
| | 2 | Olfm1 | ENSMUSG00000026833 | 1.93 | 3.35 | 1.90 | 0.79 | 0.82 | 0.00 | 0.00 |
| | 7 | Usp35 | ENSMUSG00000035713 | 11.43 | 20.10 | 9.98 | 0.81 | 1.01 | 0.00 | 0.02 |
| | 4 | Bmp8b | ENSMUSG00000002384 | 3.72 | 7.03 | 4.17 | 0.92 | 0.75 | 0.00 | 0.00 |
| | 17 | Srd5a2 | ENSMUSG00000038541 | 17.97 | 35.21 | 8.64 | 0.97 | 2.03 | 0.02 | 0.00 |
| | 9 | Cacna2d2 | ENSMUSG00000010066 | 1.79 | 3.53 | 2.24 | 0.98 | 0.66 | 0.00 | 0.05 |
| | 5 | Gm9899 | ENSMUSG00000053214 | 2.36 | 4.94 | 2.67 | 1.07 | 0.89 | 0.05 | 0.02 |
| | 18 | Galr1 | ENSMUSG00000024553 | 13.74 | 30.35 | 19.72 | 1.14 | 0.62 | 0.00 | 0.01 |
| | 18 | Mapk4 | ENSMUSG00000024558 | 1.08 | 2.43 | 1.36 | 1.17 | 0.84 | 0.00 | 0.03 |
| | 1 | Sgpp2 | ENSMUSG00000032908 | 2.74 | 6.48 | 1.12 | 1.24 | 2.53 | 0.03 | 0.00 |
| | 15 | Rpl3 | ENSMUSG00000060036 | 0.61 | 1.50 | 0.70 | 1.29 | 1.10 | 0.01 | 0.04 |
| **Placenta male** | 2 | Ltk | ENSMUSG00000027297 | 1.19 | 2.95 | 1.21 | 1.31 | 1.29 | 0.03 | 0.03 |
| | 7 | Gm26827 | ENSMUSG00000097634 | 0.39 | 0.99 | 0.58 | 1.34 | 0.77 | 0.02 | 0.01 |
| | 15 | Myc | ENSMUSG00000022346 | 0.49 | 1.26 | 0.22 | 1.36 | 2.55 | 0.04 | 0.01 |
| | 7 | Sae1 | ENSMUSG00000052833 | 0.57 | 1.48 | 0.52 | 1.39 | 1.52 | 0.00 | 0.03 |
| | 10 | Grin3b | ENSMUSG00000035745 | 4.00 | 12.16 | 5.21 | 1.60 | 1.22 | 0.00 | 0.00 |
| | 19 | Rnaseh2c | ENSMUSG00000024925 | 1.32 | 4.07 | 0.82 | 1.62 | 2.31 | 0.02 | 0.00 |
| | 6 | Cxcl12 | ENSMUSG00000061353 | 1.89 | 6.00 | 2.74 | 1.67 | 1.13 | 0.00 | 0.01 |
| | 7 | Ears2 | ENSMUSG00000030871 | 0.44 | 1.99 | 0.32 | 2.18 | 2.62 | 0.02 | 0.04 |
| | 10 | Mmp11 | ENSMUSG00000000901 | 0.78 | 6.86 | 1.82 | 3.14 | 1.91 | 0.00 | 0.00 |
| | X | Pdk3 | ENSMUSG00000035232 | 7.73 | 1.49 | 9.51 | -2.37 | -2.67 | 0.04 | 0.00 |
| | X | 3630019K06Ril | ENSMUSG00000052364 | 7.00 | 1.39 | 7.67 | -2.33 | -2.47 | 0.00 | 0.00 |
| | X | Pdzd11 | ENSMUSG00000015668 | 6.87 | 1.80 | 9.13 | -1.94 | -2.35 | 0.00 | 0.00 |
| | X | Htatsf1 | ENSMUSG00000067873 | 7.26 | 2.41 | 9.54 | -1.59 | -1.98 | 0.01 | 0.01 |
| | X | Ppp1r3f | ENSMUSG00000039556 | 5.86 | 3.22 | 9.71 | -0.86 | -1.59 | 0.04 | 0.00 |
| | X | Diaph2 | ENSMUSG00000034480 | 5.34 | 3.05 | 7.07 | -0.81 | -1.21 | 0.00 | 0.01 |
| | X | Chic1 | ENSMUSG00000031327 | 1.71 | 4.69 | 0.50 | 1.46 | 3.22 | 0.02 | 0.00 |
| | X | Gpm6b | ENSMUSG00000031342 | 4.86 | 13.44 | 0.64 | 1.47 | 4.39 | 0.00 | 0.01 |
| | X | Gm10489 | ENSMUSG00000097444 | 1.22 | 10.47 | 3.83 | 3.10 | 1.45 | 0.00 | 0.00 |
| | X | Phka2 | ENSMUSG00000031295 | 0.77 | 7.74 | 4.41 | 3.33 | 0.81 | 0.00 | 0.03 |
| | X | Timm8a1 | ENSMUSG00000048007 | 0.29 | 8.69 | 5.44 | 4.93 | 0.68 | 0.00 | 0.00 |
| | 14 | Rnf219 | ENSMUSG00000022120 | 3.49 | 0.84 | 4.87 | -2.05 | -2.53 | 0.04 | 0.04 |
| | 11 | Hoxb8 | ENSMUSG00000056648 | 13.26 | 19.41 | 11.09 | 0.55 | 0.81 | 0.00 | 0.00 |
| | 12 | Trib2 | ENSMUSG00000020601 | 0.96 | 1.52 | 1.04 | 0.67 | 0.54 | 0.00 | 0.00 |
| | 16 | Sidt1 | ENSMUSG00000022696 | 7.69 | 12.34 | 3.83 | 0.68 | 1.69 | 0.03 | 0.01 |
| | 15 | Hoxc10 | ENSMUSG00000022484 | 6.34 | 10.27 | 4.00 | 0.70 | 1.36 | 0.01 | 0.00 |
| | 5 | Pds5a | ENSMUSG00000029202 | 0.56 | 0.93 | 0.27 | 0.73 | 1.80 | 0.05 | 0.00 |
| | 10 | Shc2 | ENSMUSG00000020312 | 1.89 | 3.22 | 1.13 | 0.76 | 1.52 | 0.03 | 0.00 |
| | 9 | Tmem158 | ENSMUSG00000054871 | 0.90 | 1.55 | 0.38 | 0.79 | 2.04 | 0.04 | 0.00 |
| | 9 | Rora | ENSMUSG00000032238 | 1.12 | 3.27 | 1.13 | 0.97 | 1.53 | 0.01 | 0.00 |
| **Placenta female** | 11 | Wnt3a | ENSMUSG00000009900 | 3.73 | 7.76 | 4.44 | 1.06 | 0.81 | 0.00 | 0.01 |
| | 16 | Robo1 | ENSMUSG00000022883 | 1.04 | 4.93 | 1.41 | 2.25 | 1.80 | 0.00 | 0.00 |
| | 2 | Rims4 | ENSMUSG00000035226 | 0.65 | 3.61 | 0.81 | 2.48 | 2.16 | 0.00 | 0.04 |
| | 5 | Ttyh3 | ENSMUSG00000036565 | 1.42 | 10.14 | 1.19 | 2.84 | 3.09 | 0.00 | 0.00 |
| | 17 | Ndufb10 | ENSMUSG00000040048 | 0.48 | 8.03 | 1.67 | 4.08 | 2.27 | 0.00 | 0.00 |
| | 19 | Ighmbp2 | ENSMUSG00000024831 | 0.13 | 22.39 | 11.88 | 7.48 | 0.91 | 0.00 | 0.00 |
| | X | Syn1 | ENSMUSG00000037217 | 16.00 | 1.67 | 21.03 | -3.26 | -3.66 | 0.04 | 0.01 |
| | X | Iqsec2 | ENSMUSG00000041115 | 16.05 | 11.23 | 15.85 | -0.52 | -0.50 | 0.00 | 0.00 |
| | X | Sept6 | ENSMUSG00000050379 | 19.84 | 31.78 | 20.96 | 0.68 | 0.60 | 0.00 | 0.00 |
| | X | Usp27x | ENSMUSG00000046269 | 15.75 | 26.99 | 16.34 | 0.78 | 0.72 | 0.00 | 0.00 |
| | X | Pim2 | ENSMUSG00000031155 | 5.85 | 25.55 | 7.71 | 2.13 | 1.73 | 0.00 | 0.01 |

Chr. = Chromosome. LFC=log2 Fold Change. Cutoffs used: FDR < 0.05 in both comparisons considered as transgressive.
